# Targeting Endoplasmic Reticulum Stress and Nitroso-Redox Imbalance in Neuroendocrine Prostate Cancer: The Therapeutic Role of Nitric Oxide

**DOI:** 10.1101/2024.11.27.624202

**Authors:** Fakiha Firdaus, Osmel Companioni Napoles, Rehana Qureshi, Raul Ariel Dulce, Anusha Edupuganti, Surinder Kumar, Khushi Shah, Salih Salihoglu, Derek J Van Booven, Richard Gasca, Sanoj Punnen, David Lombard, Joshua M. Hare, Himanshu Arora

**Author notes:** These authors contributed equally (Fakiha Firdaus, Osmel Companioni Napoles, Rehana Qureshi, Himanshu Arora).

## Abstract

Neuroendocrine prostate cancer (NEPC) is an aggressive and therapy-resistant subtype of prostate cancer. Current standard-of-care treatment for NEPC involves chemotherapies, which largely exert their cytotoxic effects by forming DNA crosslinks, disrupting DNA replication and transcription in NEPC cells. However, these therapies are often met with resistance, partly due to increased endoplasmic reticulum (ER) stress, which facilitates cancer cell survival and adaptive mechanisms. Despite its critical role, the molecular landscape underlying ER stress in NEPC remains inadequately understood. Here we showed that ER stress is intimately linked to the metabolic reprogramming of NEPC cells, a process that supports their transition from adenocarcinoma to a neuroendocrine phenotype. We identified MYCN as a key driver of this process, promoting unfolded protein response (UPR) elements that enhance ER stress by increasing the efflux of calcium ions through the ER which later is absorbed by the mitochondria and assist in increasing the overall glycolytic stress, thereby adding to the extended survival and metastatic potential of an NEPC cell. Our previous studies highlighted the importance of S-nitrosylation as a protein modification that is dysregulated in high-grade PCa. In this context, structural analysis of MYCN revealed potential S-nitrosylation sites at the positions Cys4, 186, and 464, respectively. However, similar to the castration-resistant stage, this modification is hindered in NEPC due to impaired nitric oxide (NO) production from dysregulated endothelial nitric oxide synthases (eNOS). We found that exogenous NO supplementation S-nitrosylates MYCN, reducing its binding to protein molecules which are essential to assist with increasing ER stress in NEPC cells. Exogenous supplementation of NO reduced the overall tumor burden in the mice harboring orthotopic NEPC cells and reduced the metastasis to the brain and liver. In conclusion, the findings from this study enrich our understanding of the mechanisms driving the ER stress responses in NEPC phenotype and how NO supplementation could pave the way as potential therapeutics for this challenging cancer.

**HIGHLIGHTS:**

- Endoplasmic reticulum (ER) stress is intricately linked to metabolic reprogramming, which supports the transition from prostate adenocarcinoma to neuroendocrine prostate cancer (NEPC).
- MYCN increases the ER stress in NEPC cells and is correlated with increased nitroso-redox imbalance.
- Structural analysis reveals potential S-nitrosylation sites on MYCN. Exogenous nitric oxide (NO) supplementation induces S-nitrosylation, disrupting MYCN’s role in enhancing ER stress.
- NO supplementation reduced tumor burden and metastasis in NEPC-bearing mice, highlighting its potential as a therapeutic avenue for NEPC.
- Exogenous NO supplementation inhibits ER stress by targeting unfolded protein response (UPR) elements and decreasing calcium ion efflux, inhibiting the glycolytic stress in NEPC.

**Graphical Abstract:** 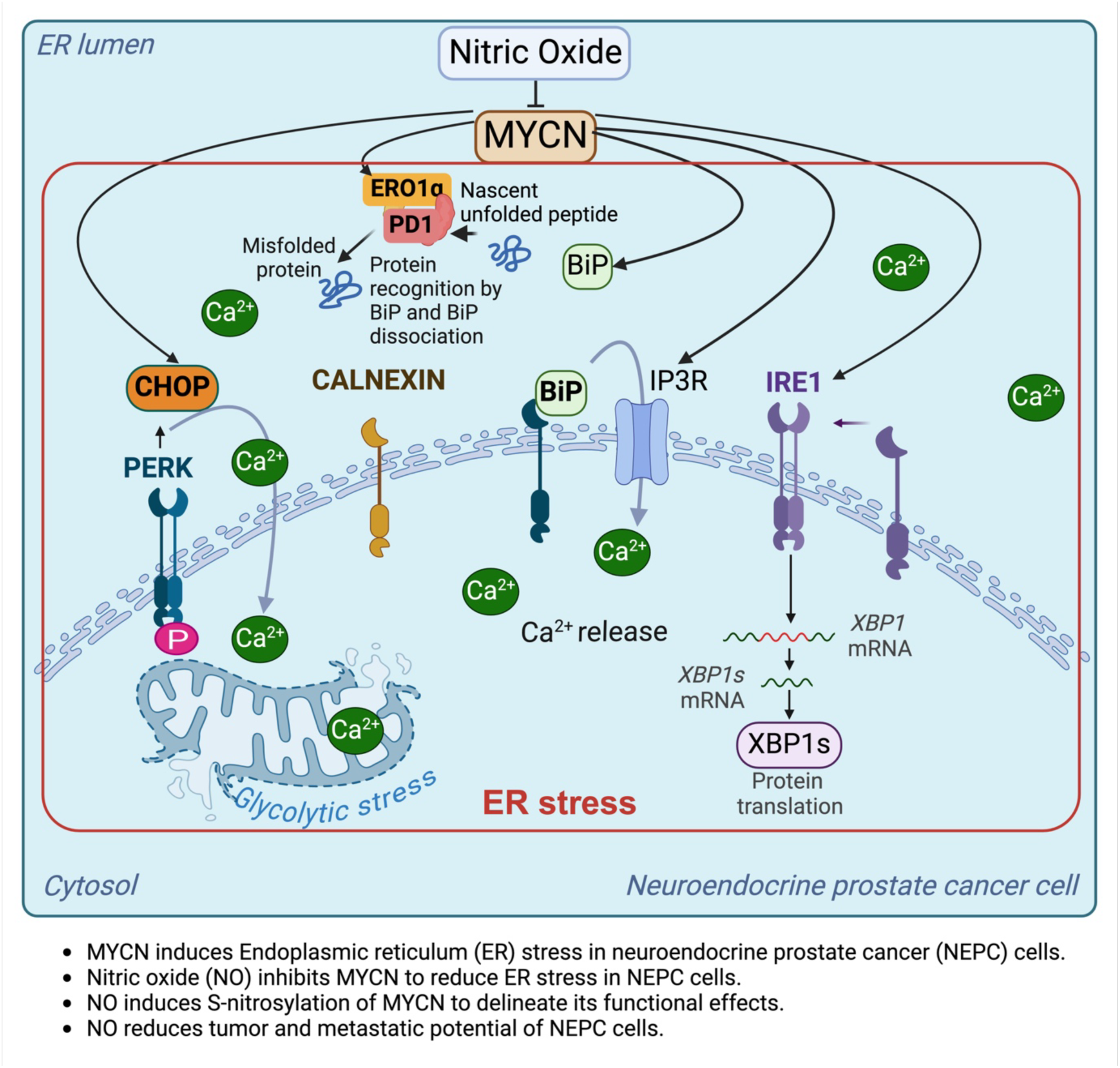

## INTRODUCTION

Neuroendocrine prostate cancer (NEPC) is an aggressive and highly therapy-resistant subtype of prostate cancer, contributing to up to 25% of prostate cancer-related deaths ^1,2^. NEPC typically arises as an adaptive response to androgen deprivation therapies (ADT) in late-stage, castration-resistant prostate cancer (CRPC), a process that is often marked by transdifferentiation from an adenocarcinoma to a neuroendocrine phenotype ^3,4^. This transformation, known as lineage plasticity, enables cancer cells to evade conventional therapies, such as androgen receptor (AR)-targeted therapies and taxane-based chemotherapies, rendering them highly resistant and contributing to rapid disease progression ^5,6^. Current treatment strategies for NEPC, including chemotherapies in combination with taxanes or etoposide, have shown limited success in improving patient outcomes ^5,6^. Ongoing clinical trials exploring combinations of chemo-immunotherapies, such as nivolumab with cabazitaxel and carboplatin, have yet to demonstrate significantly enhanced efficacy ^5,6^. A key factor driving NEPC progression is the ability of cancer cells to tolerate severe metabolic and environmental stressors through endoplasmic reticulum (ER) stress adaptation ^7,8^. The ER stress response, or the unfolded protein response (UPR), is activated by the accumulation of misfolded or unfolded proteins in the ER, triggering signaling pathways aimed at restoring cellular homeostasis ^7,8^. However, prolonged or unresolved ER stress can lead to mitochondrial dysfunction, oxidative stress, and activation of pro-survival pathways, all of which promote tumor growth, therapy resistance, and metastasis ^9–12^. In NEPC, heightened ER stress not only supports cell survival under metabolic duress but also promotes the transition from an androgen-dependent adenocarcinoma to an androgen receptor (AR)-independent neuroendocrine phenotype ^13–16^. This transition is tightly regulated by oncogenic drivers such as MYCN, which play a pivotal role in reprogramming gene expression and promoting metabolic reorganization to meet the increased biosynthetic demands of aggressive cancer cells ^13,17^.

MYCN is particularly important in NEPC, where its overexpression drives epigenetic reprogramming, enhances UPR activation, and fosters lineage plasticity ^13,17^. Recent studies have demonstrated that MYCN promotes ER stress by increasing calcium ion efflux from the ER, which is then absorbed by mitochondria, further increasing glycolytic and oxidative stress ^13,17^. These stress adaptations create a feedback loop that supports the aggressive behavior of cancer cells, contributing to therapy resistance and metastatic potential. Notably, our previous studies have shown that S-nitrosylation, a post-translational modification that involves the covalent attachment of a nitric oxide (NO) group to cysteine residues, plays a critical role in regulating protein function and modulating cancer cell signaling ^18^. However, in high-grade PCa, the production of NO is significantly impaired due to dysregulation in endothelial nitric oxide synthase (eNOS) activity ^18^. This may prevent the S-nitrosylation of MYCN and contribute to the persistence of ER stress.

Given the central role of ER stress and MYCN in driving NEPC progression, we hypothesized that exogenous supplementation of NO could restore the S-nitrosylation of MYCN, thereby reducing ER stress and mitigating the aggressive behavior of NEPC cells. Nitric oxide has been shown to have a range of cytoprotective effects, including reducing oxidative stress and restoring normal cellular homeostasis. By reinstating the S-nitrosylation of MYCN, NO-based therapies could re-regulate key signaling pathways that promote ER stress, inhibit transdifferentiation, and reduce the overall tumor burden in NEPC.

In this study, we explored the molecular mechanisms by which ER stress and MYCN contribute to the pathogenesis of NEPC and investigated the therapeutic potential of NO supplementation in re-regulating these processes. Using a combination of *in vitro* cell models and an in-vivo orthotopic murine xenograft model, we demonstrated that NO treatment effectively restored MYCN S-nitrosylation, reduced ER stress markers, and inhibited the transdifferentiation of prostate adenocarcinoma cells to neuroendocrine phenotype. Moreover, we observed a significant reduction in tumor growth and metastasis following NO treatment, highlighting the potential of NO-based therapies as a novel strategy to combat therapy-resistant NEPC.

Together, our findings provide critical insights into the molecular mechanisms underlying NEPC progression and suggest that NO supplementation represents a promising therapeutic avenue for targeting ER stress and lineage plasticity in this aggressive cancer subtype.

## RESULTS

### ER Stress Biomarkers Correlate with Prostate Cancer Progression and Survival Outcomes

To investigate the relationship between ER stress biomarkers and PCa progression, RNA sequencing (RNAseq) data from 500 patients in the Cancer Genome Atlas (TCGA) prostate adenocarcinoma (PRAD) cohort and 4,983 patients from the Decipher GRID database were analyzed. Although the Decipher GRID and TCGA databases primarily consist of PRAD samples, they remain valuable for studying the molecular landscape relevant to NEPC, considering the majority of the cases of NEPC evolve from adenocarcinoma, especially therapy-induced NEPC (t-NEPC), following androgen deprivation therapy (ADT) ^19,20^. Therefore, the patients were stratified based on their Gleason scores (GS) and clinical parameters such as age, PSA levels, and therapy history. In the TCGA cohort, the distribution included 52 noncancerous controls, 45 patients with GS6, 247 with GS7, 64 with GS8, 136 with GS9, and 4 with GS10. The Decipher GRID cohort comprised 382 patients with GS6, 3,544 with GS7, 446 with GS8, and 607 with GS9. The differential expression analysis revealed a significant upregulation of several key ER stress markers—IRE1 (ERN1), CANX (Calnexin), CHOP (DDIT3), XBP1, and PDI (protein disulfide isomerase-A1)(Figure 1A, Supplementary Figure 1) in high-grade tumors (GS ≥ 8), compared to lower-grade tumors and noncancerous controls (Figure 1A-B). This upregulation was consistently observed across both the TCGA and Decipher GRID cohorts, suggesting that ER stress is a prominent feature of advanced PCa.

**Figure 1:**
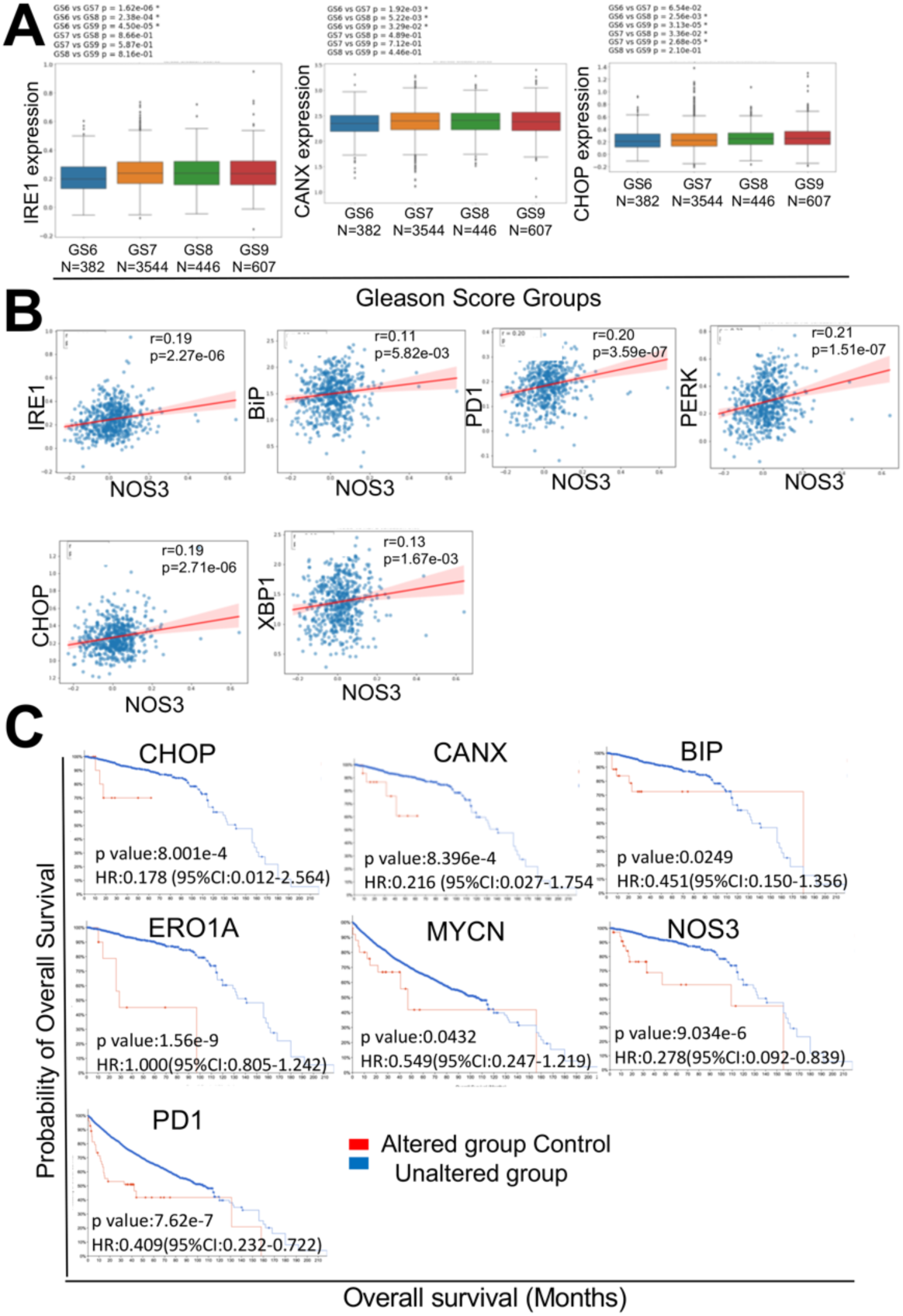
An enrichment analysis showing ER stress biomarkers correlate with prostate cancer (PCa) progression and survival outcomes. **(a)** The expression of ER stress markers in PCa patients was obtained from the Decipher Grid. The box plots span the 25th to 75th percentiles, and the expression is represented as log⁡10log10 (reads per kilobase million, RPKM). The median (midline of box plots) and maximum and minimum (upper and lower bounds of whiskers) values are depicted. Significance was determined by a non-parametric Wilcoxon signed-rank test (two-sided), and P*P* values correspond to the comparison between Gleason score groups with other subgroups of PCa patients. **(b)** The plots show a significant positive correlation between ER stress markers (IRE1, BiP, PDI, PERK, CHOP, and XBP1) with NOS3 in the Decipher Grid data. **(c)** Kaplan–Meier survival curves segregated by BiP, CHOP, CANX, ERO1A, MYCN, NOS3, and PDI expression. Overall survival is significantly diminished for individuals with high expression. *P* values were calculated using a log-rank (Mantel–Cox) test.

To explore the link between nitric oxide signaling and ER stress, we performed a correlation analysis between NOS3 (endothelial nitric oxide synthase) and ER stress markers. A significant positive correlation was observed between NOS3 and IRE1 (r=0.19, p=2.27e-06), BiP (r=0.11, p=5.82e-03), PDI (r=0.20, p=3.59e-07), XBP1 (r=0.13, p=1.67e-03), PERK (r=0.21, p=1.51e-07), and CHOP (r=0.19, p=2.71e-06) in the Decipher GRID dataset (Figure 1B). These correlations suggest that NOS3 dysregulation is associated with increased ER stress in high-grade PCa, potentially contributing to tumor progression by amplifying the stress response.

To assess the clinical significance of ER stress markers in PCa, we conducted a survival analysis using the cBioPortal platform. Kaplan-Meier survival curves showed that high expression of BiP, CHOP, and IRE1 was associated with significantly reduced overall survival in PCa patients (Figure 1C). For example, elevated CHOP expression was associated with a hazard ratio (HR) of 0.178 (p=8.001e-4), while high CANX (Calnexin) expression yielded an HR of 0.216 (p=8.396e-4). Additionally, increased PDI expression was correlated with poor survival outcomes (HR=0.409, p=7.62e-7). These findings suggest that elevated ER stress marker expression is linked to worse clinical outcomes, particularly in high-grade PCa patients.

### MYC increases the ER stress responses in PCa cells and these responses are correlated with increased nitroso-redox imbalance

We developed test models of AR loss using MyC-CaP cells, an epithelial-like cell line that was originally developed from the prostate of a 16-month-old, male mouse with PCa. For this, MyC-CaP cells were subjected to the partial loss of AR (MyC-CaP^ShAR^) or the complete loss (MyC-CaP^APIPC^) using SMARTvector Inducible Lentiviral shRNA vector. Steps involved in the generation of MyC-CaP^APIPC^ from MyC-CaP^shAR^ are highlighted in the Supplementary Figure 2A). MyC-CaP^APIPC^ cells showed complete AR absence (Figure 2A) and increased MYCN expression, which suggests neuroendocrine phenotype formation (Supplementary Figure 2B). To elucidate the relationship between nitroso redox imbalance and ER stress and to assess the involvement of mitochondrial dysfunction in this interplay, we used LNCaP, 22Rv1, H660, MyCAP, MyC-CaP^shAR^ and MyC-CaP^APIPC^ cells. The mitochondrial mass was quantified using fluorescent mitochondrial dyes, which provided insights into the bioenergetic health of the cells. We observed a significant increase in mitochondrial mass in cells representing high-grade PCa (Figure 1B). The unfolded protein response (UPR) markers, such as IRE1^21^, BIP^22^, Calnexin ^23^, CHOP ^24^, Ero1-La ^25^, and PERK ^26^ were examined to gauge ER stress levels using LNCaP and H660 cells. We observed a notable upregulation in these markers, particularly in H660 cells, indicating a heightened ER stress response in NEPC (Figure 1C).

**FIGURE 2:**
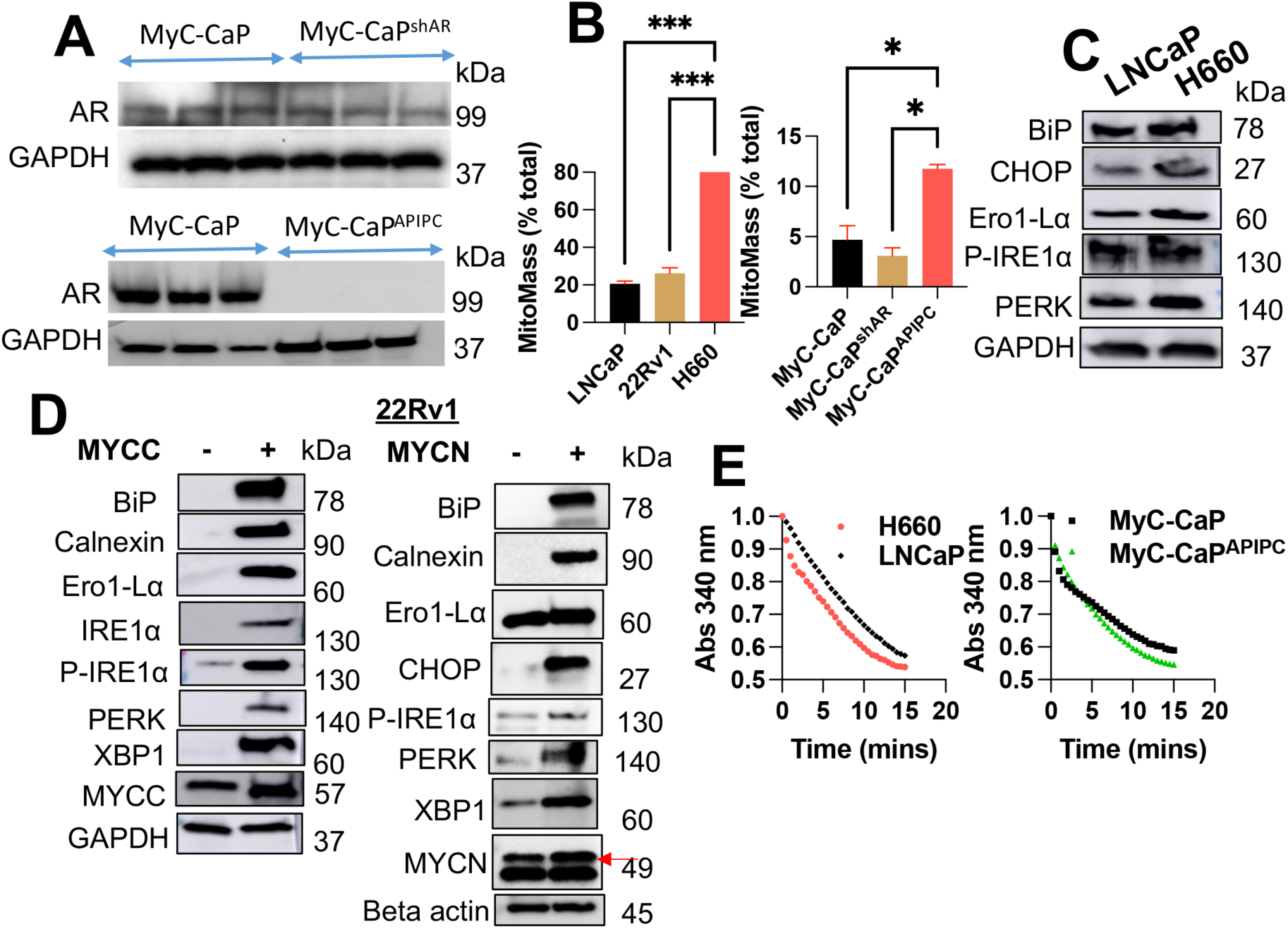
MYC Enhances ER Stress Responses and Nitroso-Redox Imbalance in PCa Cells. **(a)** Western blot analysis showing androgen receptor (AR) levels in MyCaP, MyCaPshAR, and MyCaPAPIPC cell lines. GAPDH serves as a loading control. **(b)** Bar graph depicting percent Mitomass in various PCa cell lines (LNCaP, 22Rv1, H660, MyCaP, MyCaPshAR, and MyCaPAPIPC). **(c)** Western blot comparing ER stress markers between LNCaP and H660 cell lines. GAPDH serves as a loading control. **(d)** Western blot showing ER stress marker expression in 22Rv1 cells with and without stable MYCC or MYCN overexpression. GAPDH serves as a loading control. (**e)** Bar graphs comparing GSNOR activity between LNCaP and H660 cells (left panel) and between MyCaP and MyCaPAPIPC cells (right panel). Data represent mean ± standard deviation from three independent biological replicates (p < 0.001, two-way ANOVA).

Further, we measured nitrate concentrations in these cell lines to estimate nitrosative stress (Supplementary Figure 3). A negative correlation was established between nitrate concentrations (which were reduced) and the expression of Cysteine Sulfenic Acid (CSA) (which was increased) in our earlier study ^18^, underscoring the interconnection between nitrosative stress and hypoxic conditions in PCa. Next, to determine the impact of MYC on ER stress response elements, we generated 22Rv1 cells that stably overexpressed MYCC and MYCN. The rationale for over-expressing MYCC along with MYCN in 22Rv1 cells is the fact that CRPC cells do not biologically express MYCN, and doing so may not represent the typical stress responses seen in CRPC^27,28^. Subsequent analyses revealed an exacerbation of ER stress markers, including BiP, Calnexin, Ero1-Lα, IRE1α, p-IRE1α, PERK, and XBP1, upon both MYCC and MYCN overexpression (Figure 2D). These findings provide direct evidence of MYCN role in amplifying the malignant phenotype through the induction of ER stress.

Next, we confirmed that similar to CRPC^18^, in NEPC cells (H660), diminished tetrahydrobiopterin to the 7,8-dihydrobiopterin ratio (BH4:BH2) ratio led to the destabilization of NOS subunits (uncoupling) and shift the enzymatic activity towards superoxide (O2-) generation ^29–33^. This is reflected by a subsequent decrease in the activity of S-Nitrosoglutathione reductase (GSNOR) in proteins derived from H660 and MyC-CaP^APIPC^ cells compared to LNCAP and MyC-CaP cells (Figure 2E). This decrease in GSNOR activity in high-grade PCa cells implies an accelerated depletion rate of S-Nitrosoglutathione (GSNO) in NEPC ^18^. Together, these results conclusively demonstrate that MYC increases ER stress, and these responses are correlated with increased nitroso-redox imbalance.

### Nitric oxide supplementation could inhibit MYC and overcome ER stress responses in PCa cells

Under conditions of ER stress, the ER’s ability to regulate calcium homeostasis is compromised in neuroendocrine cells ^34,35^. As a result, abundant calcium ions could be released into the cytoplasm through channels such as inositol 1,4,5-triphosphate receptors (IP3Rs) and ryanodine receptors (RyRs). These elevated Ca^+2^ are taken up by the mitochondria, and excessive intake can further enhance the production of ROSs, which exacerbates oxidative stress and, therefore, adds to the mitochondrial dysfunction ^36^. To confirm if MYC has a direct effect on Ca^+2^ release in high-grade PCa cells, we evaluated Ca^+2^ in 22Rv1 cells, which overexpresses MYCC and MYCN. The cells were loaded with FURA-2, fluorescence was acquired, and ratiometric data were analyzed. Results showed that the 22Rv1 overexpressing MYCN showed a significant increase in the Ca^+2^ release comparable to the MYCC overexpressing or control (CaCl_2_ group) (Supplementary Figure 4). Previous studies from our group have elucidated NO’s antioxidative properties in PCa cell lines [45]. Expanding on this foundation, we aimed to explore whether NO supplementation (using S-nitroso glutathione, an NO donor) could inhibit ER stress responses. In this context, first, we tested the inhibitory effects (if any) of NO treatment on reducing ER stress-induced Ca^+2^ release. For this, we used LNCaP, 22Rv1, and H660 cells treated with either vehicle or GSNO (50uM), which were loaded with FURA-2. Subsequently, the fluorescence was acquired, and ratiometric data were analyzed. Results showed that the GSNO decreased the Ca^+2^ release in CRPC (22Rv1) and NEPC (H660) cells and not in primary PCa cells (LNCaP) comparable to the positive control (CaCl_2_ group) (Figure 3A).

**FIGURE 3:**
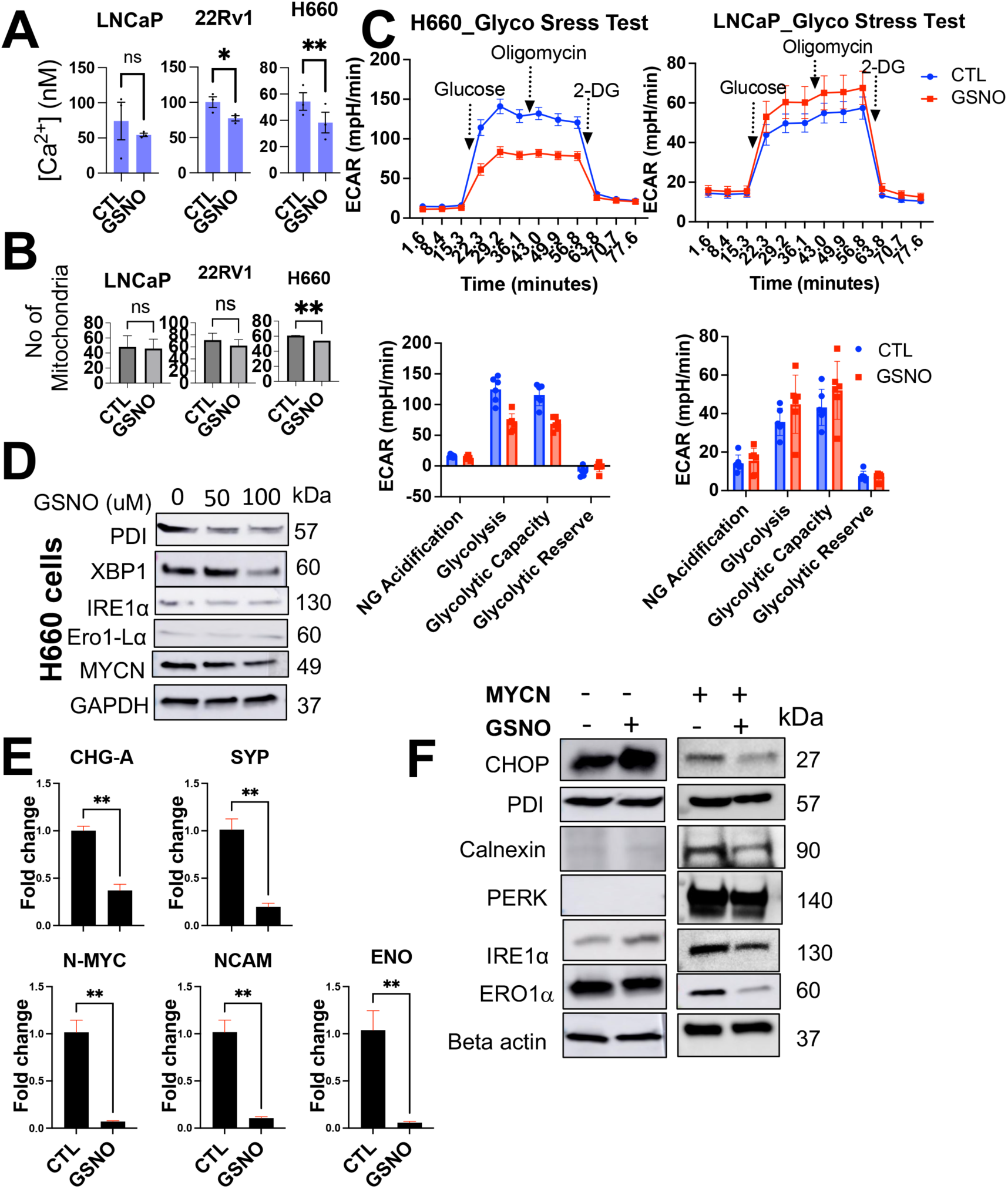
Nitric Oxide Supplementation Inhibits MYC and Inhibits ER Stress Responses in Prostate Cancer Cells. **(a)** Bar graph showing overall calcium levels in LNCaP, 22Rv1, and H660 cells with or without GSNO treatment (50 μM). **(b)** Bar graph depicting the number of mitochondria in LNCaP, 22Rv1, and H660 cells with or without GSNO treatment (50 μM), assessed using mitochondrial dye. **(c)** Results of Seahorse XF Mito Stress Tests on LNCaP and H660 cells with or without GSNO treatment. The graph shows a double plot of Extracellular Acidification Rate (ECAR) from three independent biological replicates. **(d)** Western blot analysis showing the inhibitory effects of GSNO treatment on ER stress markers in H660 cells. GAPDH serves as a loading control. **(e)** Quantitative real-time PCR results demonstrating the inhibitory effects of GSNO treatment on neuroendocrine prostate cancer (NEPC) markers in H660 cells. Data represent mean ± standard deviation from three independent biological replicates (p < 0.001). **(f)** Western blot analysis showing the inhibitory effects of GSNO treatment on ER stress markers in MYCN overexpressing 22Rv1 cells. GAPDH serves as a loading control.

To further investigate NO’s downstream bioenergetic and metabolic impacts (if any) on NEPC cells, we treated LNCaP, 22Rv1, and H660 cells with GSNO (50 µM) over a 48-hour period. Post-treatment, we evaluated the number of mitochondria in the cells (an indicator of cellular health and bioenergetics) using quantitative fluorescence microscopy, employing MitoTracker Green FM. Notably, we observed a significant decrease in the number of mitochondria in H660 cells (Figure 3B). Similar patterns were observed when comparing MyCAP, MyC-CaP^shAR^ and MyC-CaP^APIPC^ cells (Supplementary Figure 5). Additionally, we examined whether the observed decrease in mitochondrial count corresponded with changes in mitochondrial function. For this, we investigate the bioenergetic and metabolic impact of NO through Seahorse XF Mito Stress Tests on LNCaP and H660 cells. The cells were treated with 50 µM of GSNO for 48 hours before subjecting them to the Seahorse test. This assay included sequential injections of 2 µM Oligomycin (to inhibit ATP synthase (complex V)) and 2 µM FCCP (Carbonyl cyanide 4-(trifluoromethoxy) phenylhydrazone) (to uncouple oxidative phosphorylation by dissipating the proton gradient, allowing for the measurement of maximal respiration) to measure mitochondrial function and the oxygen consumption rate (OCR). Results highlighted a significant reduction in glycolysis and glycolytic capacity in H660 cells alone. No significant impact was observed on mitochondrial respiration (basal respiration, maximal respiration, proton leak, ATP production, and spare respiratory capacity). A slight reduction was observed in non-mitochondrial oxygen consumption rate in H660 cells post-NO treatment compared to LNCaP cells (Figure 3C).

To further study the effects of NO supplementation against the ER stress at molecular levels, we monitored the expression of IRE1α, Ero1-Lα, PDI, and XBP-1, respectively, post-treatment of H660 cells with GSNO at 50 and 100uM concentrations. These UPR markers, displayed decreased expression levels upon GSNO treatment (Figure 3D). Additionally, the expression of Chromogranin (CHG-A), Synaptophysin (SYP), MYCN, Neural cell adhesion molecule (NCAM) and Enolase (ENO) was diminished in the presence of NO in H660 cells (Figure 3E). This observation suggests a downregulation of pathways associated with aggressive tumor behavior and hypoxic adaptation ^37,38^, pointing to NO’s capacity to downregulate oncogenic signaling cascades. Furthermore, we subjected the MYCN overexpressing 22Rv1 cells to GSNO to determine if NO supplementation could overcome MYCN-induced ER stress response in high-grade PCa cells. Cells with enforced MYCN expression coupled with NO treatment exhibited reduced ER stress markers relative to their counterparts overexpressing MYCN without NO treatment (Figure 3F). This delineates the inhibitory role of NO against the deleterious mitochondrial effects induced by MYCN overexpression. Together, these results underscore the potential therapeutic application of NO in attenuating MYCN-driven ER stress in high-grade PCa.

To further elucidate the molecular landscape of NEPC cells and assess the effects of NO supplementation in greater detail, we treated H660 cells with 50 µM GSNO for 48 hours, followed by RNA isolation and subsequent RNA sequencing (Figure 4A-C). The enrichment analysis uncovered a significant downregulation of pathways associated with ER stress signaling, including genes such as EXT1, TRPS2, CHDS, CDC42, and RAC ^39^ (Figure 4D, right). These pathways are integral to ER stress responses, and their downregulation indicates a reduction in the accumulation of misfolded proteins, which in turn enhances cellular homeostasis. This suggests that NO supplementation inhibits ER stress, likely by promoting the proper folding and processing of proteins within the ER.

**FIGURE 4:**
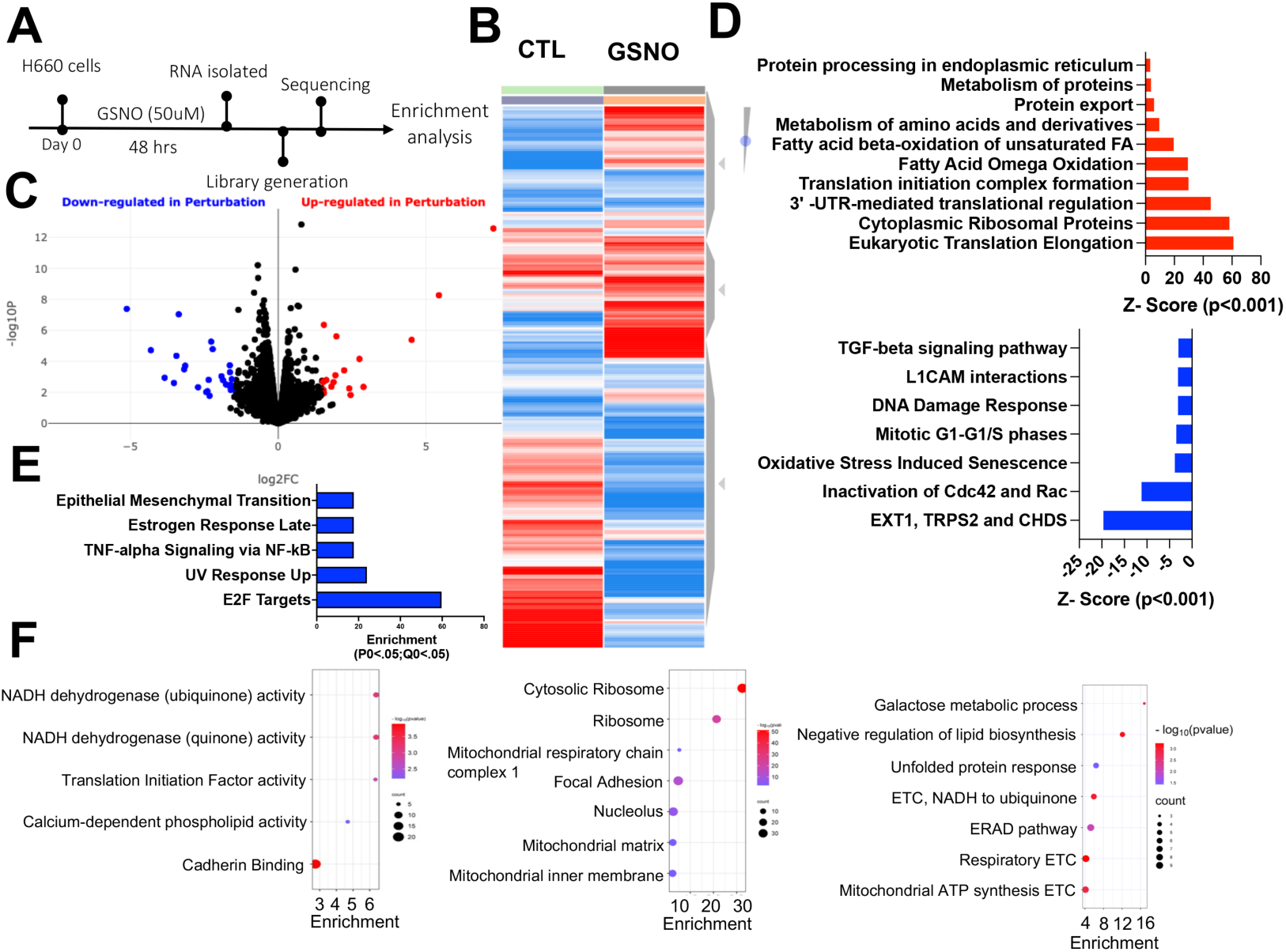
RNA Sequencing Reveals GSNO-Induced Transcriptional Changes in H660 Cells. **(a)** Schematic illustrating the RNA sequencing workflow for H660 cells treated with 50 μM GSNO for 48 hours. **(b)** Heatmap depicting significantly differentially expressed genes (fold change > 2, q-value < 0.05) following 48-hour GSNO treatment. **(c**) Visualization of differentially expressed genes (DEGs) with fold change > 1.5 and adjusted P < 0.05. Up-regulated genes are shown in red, down-regulated genes in blue. **(d**) Bar graph highlighting key downregulated pathways, including ER stress signaling (featuring genes EXT1, TRPS2, CHDS, CDC42, and RAC), oxidative stress, cell cycle regulation, DNA damage response, L1CAM interactions, and TGF-beta signaling. **(e)** Bubble plot illustrating upregulated molecular processes. **(f)** Bubble plot showing enriched cellular processes. **(g)** Bubble plot depicting upregulated biological processes, including protein synthesis and metabolic pathways such as eukaryotic translation elongation, ribosomal protein synthesis, protein processing in the ER, fatty acid oxidation, and amino acid metabolism.

Additionally, key oxidative stress pathways linked to cellular senescence were markedly reduced, consistent with decreased oxidative damage and improved stress resilience. These pathways are central to the progression of neuroendocrine differentiation and cancer aggressiveness. Further downregulated pathways included those involved in G1/G1-S cell cycle regulation, DNA damage response mechanisms ^40^, L1CAM interactions ^41^, and TGF-beta signaling ^42^ (Figure 4D), pointing to a broad suppression of stress and proliferation-related pathways in NEPC cells. Simultaneously, pathways related to protein synthesis and metabolism were significantly upregulated following GSNO treatment (Figure 4D). These included critical processes such as eukaryotic translation elongation, ribosomal protein synthesis, and 3’-UTR-mediated translational regulation ^43^, suggesting an enhanced capacity for protein synthesis. Furthermore, the initiation of translation complex formation ^44^, protein export ^45^, and protein processing in the ER ^26^ were also upregulated, indicating improved efficiency in managing ER load and preventing stress accumulation. This improvement in protein synthesis machinery suggests a more robust cellular response to meet the demands of protein folding and processing.

Moreover, Figure 4E shows the pathways related to metabolic processes were enriched, with notable upregulation in fatty acid omega oxidation ^46^, mitochondrial fatty acid beta-oxidation of unsaturated fatty acids, and amino acid metabolism ^46^. These enhancements reflect a shift toward greater metabolic activity, increased energy production, and lipid homeostasis. The improved balance in lipid metabolism would support membrane integrity and functionality in the ER, further contributing to the reduction in ER stress. Enhanced energy production, coupled with the increased availability of molecular building blocks for protein synthesis, suggests that GSNO supplementation supports a more efficient cellular environment capable of maintaining protein homeostasis. Together, these findings indicate that GSNO significantly modulates both ER stress and metabolic pathways in NEPC cells. The downregulation of stress-associated pathways and the simultaneous upregulation of protein synthesis and metabolism-related pathways suggest that NO supplementation via GSNO enhances cellular resilience, promotes protein homeostasis, and may attenuate the malignant progression of NEPC through these mechanisms.

### Exogenous induction of NO treatment demonstrates an inhibitory role against NEPC functions

Next, we studied how NO treatment could disrupt the functional properties of high grade PCa cells. For this, we treated PC3, DU145, and H660 cells, with GSNO. Our results highlighted a suppression of colony formation in all the cell lines following NO treatment (Figure 5A). In a similar context, MTT assays confirmed a significant inhibition of the proliferative capacity associated with NEPC (Figure 5B). Furthermore, we evaluated the effects of GSNO treatment on tumor-like growth in 3D culture systems. PC3, DU145, and H660 cells were cultured in 3D conditions, which more closely mimic the in vivo tumor microenvironment. GSNO treatment at a concentration of 50 μM for 9 days significantly decreased tumoroid formation across all three cell lines, with the strongest effects in H660 cells (Figure 5C). This reduction in tumoroid size and number suggests that NO signaling exerts inhibitory effects on PCa growth in a 3D environment, further highlighting its therapeutic potential.

**FIGURE 5.**
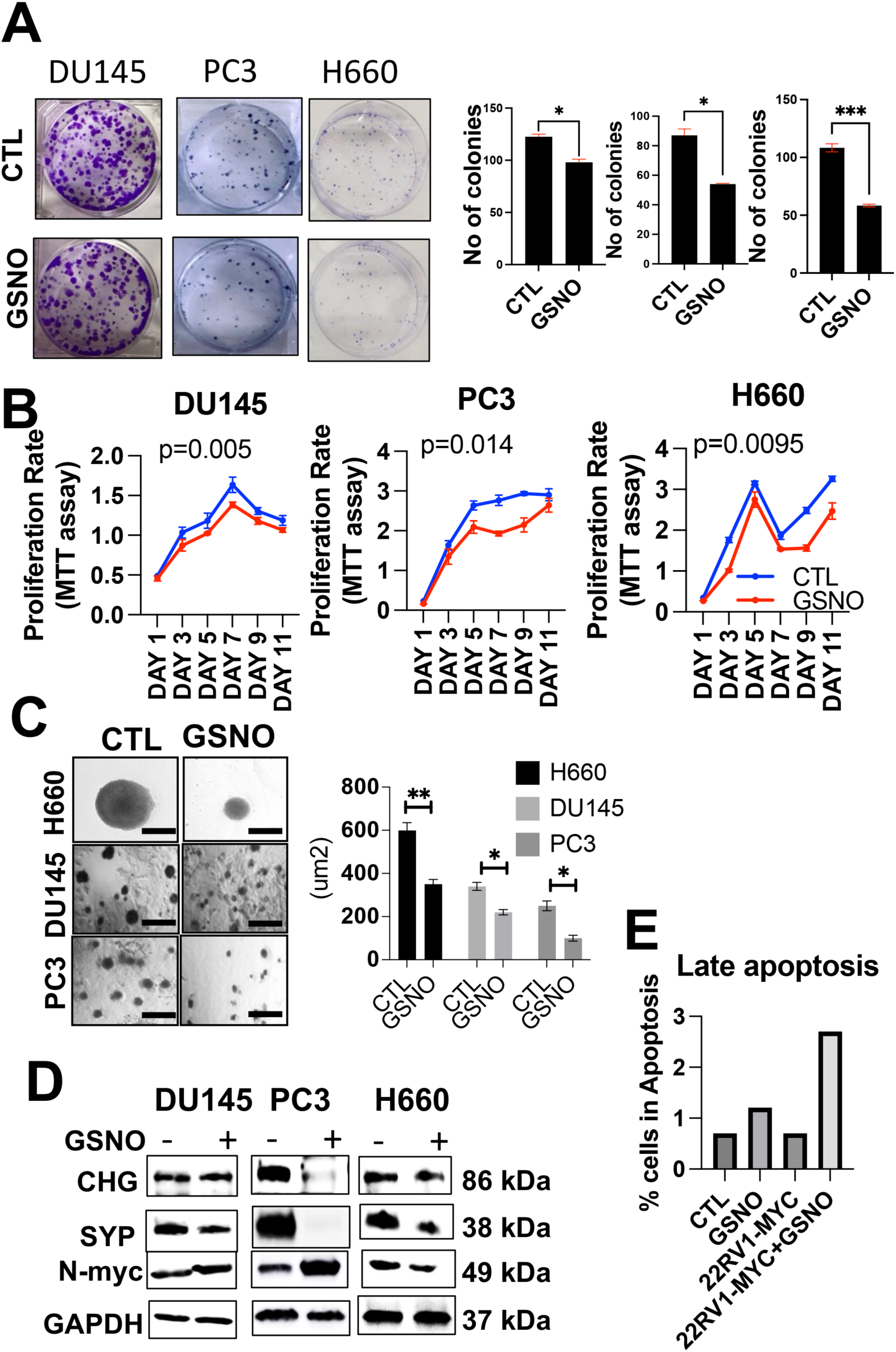
Exogenous induction of NO treatment demonstrates an inhibitory role against. **(a)** Bar graph showing the inhibitory effects of GSNO treatment on the colony-forming capabilities of DU145, PC3, and H660 cells. (**b)** Line graph illustrating the inhibitory effects of GSNO on cell proliferation (MTT assay) in DU145, PC3, and H660 cells over time. **(c)** effects of GSNO (50 μM) treatment on H660, DU145, and PC3 3D tumoroids upon 9 days of treatment **(d)** Western blot results show neuroendocrine markers such as chromogranin, synaptophysin, and MYCN levels in PC3, DU145, and H660 protein lysates before and after GSNO treatment. **(e)** Bar graph demonstrating the pro-apoptotic effects of GSNO treatment on H660 cells and 22Rv1 cells overexpressing MYC. Data represent mean ± standard deviation from three independent biological replicates (p < 0.001, two-way ANOVA).

Next, we assessed the expression of NEPC markers, such as MYCN, CHG-A, SYP, and ER stress markers, such as PERK, CANX, BIP, and Ero1-L, to quantify the treatment’s effectiveness in DU145, PC3 and H660 cells. Notably, H660 cells exhibited a marked downregulation of these markers, while PC3 cells followed a similar yet less pronounced trend. The DU145 cell line showed a moderate response, indicating an NEPC phenotype-specific sensitivity to NO (Figure 5D, Supplementary Figure 6A).

Moreover, we checked for the effects of NO in inducing apoptosis in H660 cells and in 22Rv1 cells, which stably overexpress MYC. The results highlighted an increase in overall late apoptosis upon NO supplementation in both conditions (Figure 5E). The concurrent decline in NEPC marker expression, inhibitory effects on colony formation, and increase in late apoptosis establishes the disruptive role of exogenous NO on NEPC functions. Complementing this, we investigated the NO-mediated inhibitory impact on the transition dynamics of PCa, specifically from AR-dependent to AR-independent NEPC stages. For this, we subjected MyC-CaP, MyC-CaP^shAR,^ and MyC-CaP^APIPC^ cells to similar NO treatment. The MyC-CaP^APIPC^ cells displayed reduced colony-forming and cell proliferating capabilities (Supplementary Figure 6B-C), reinforcing the potential of NO as an inhibitor of NEPC progression. Together, the results reveal that exogenous induction of NO treatment possesses a discernible inhibitory role against the functional properties of NEPC cells. The suppression of proliferative capacity and downregulation of NEPC-specific markers provide compelling evidence of NO’s therapeutic promise.

### Exogenous induction of NO demonstrates tumor inhibitory role against NEPC

To assess the impact of NO on NEPC tumor burden, we developed an orthotopic murine xenograft model using H660 cells ^47^. The cells were transduced with CMV-Firefly luciferase lentivirus and selected using puromycin at a concentration of 2 ug/ml. Amplified cells were then orthotopically injected into the dorsal prostate of mice, as described by Shahryari et al ^48^. Each mouse received an injection of 3 million NEPC cells. Following injection, the mice were imaged using IVIS after luciferin injection (i.p.), with weekly imaging performed for four weeks to monitor tumor growth. GSNO was administered at 10 mg/kg/day intraperitoneally for the duration of the study. At the end of week 4, the mice were sacrificed, and tumor volumes were estimated using mean fluorescent intensity (MFI) (Figure 6A). The results indicated that GSNO treatment significantly reduced tumor growth (*p ≤ 0.05) compared to untreated control mice, as shown in Figure 6B-C.

**Figure 6:**
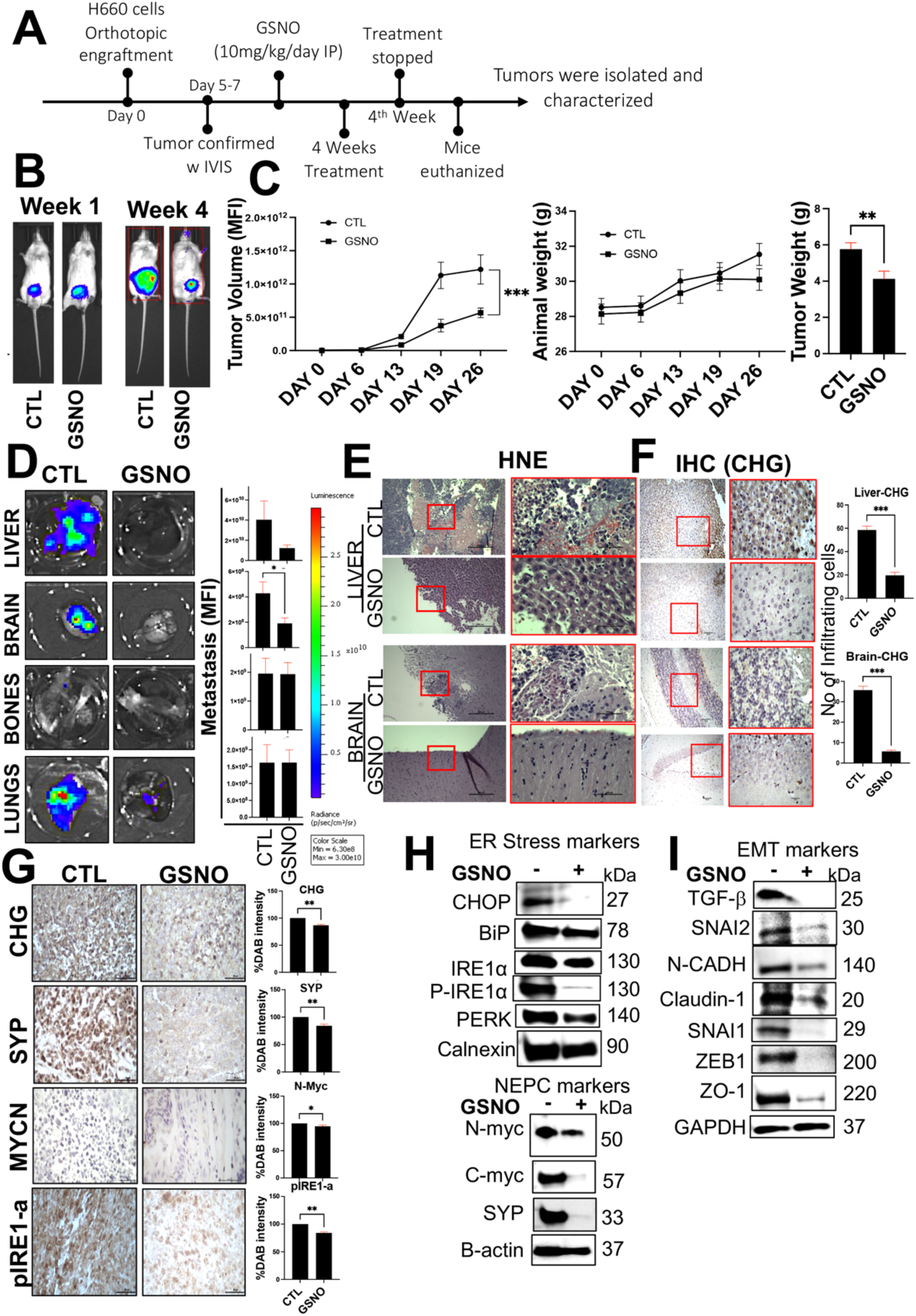
Exogenous Nitric Oxide Inhibits NEPC Tumor Growth and Metastasis: **(a)** Schematic illustrating the protocol used to evaluate the effects of NO stimulation on NEPC tumor burden. **(b)** Left: Representative bioluminescence imaging (BLI) of NOD-SCID mice (n=8) orthotopically injected with H660-Luciferase tagged cells at week 0 and week 4. Right: Line graph showing mean tumor volumes over time for GSNO-treated and control groups. **(c**) Bar graphs depicting: Tumor volumes, Animal weights, Tumor weights in mice treated with or without GSNO (10 mg/kg/day, intraperitoneal). **(d**) Ex-vivo Imaging of Metastases: Left: Representative ex-vivo bioluminescence images of liver, brain, bone, and lung tissues highlighting metastatic areas. Right: Bar graph showing mean normalized photon flux/second (±SEM) from metastases in each tissue type. Student’s t-test, **p < 0.001. (**e)** Hematoxylin and eosin (H&E)-stained sections of liver and brain tissues. Arrows indicate metastatic lesions. **(f)** Images showing immunohistochemical staining for chromogranin (CHG) neuroendocrine markers in liver and brain tissue sections. **(g)** Images showing immunohistochemical staining for pIRE1-α, synaptophysin (SYP), and chromogranin A (CHG) in tumor sections from each group. Scale bar: 20 μm. **(h)** Western blot results showing NEPC and ER stress markers levels in tumor lysates from GSNO-treated and control mice. **(i)** Western blot results showing levels of EMT markers in tumor lysates from GSNO-treated and control mice.

Additionally, GSNO treatment resulted in a significant reduction in metastatic colonization to the brain and liver as assessed by bioluminescence imaging (BLI) and metastatic incidence (Figure 6D). No metastasis was found in bones, and no significant reduction in lung metastasis (Figure 6D-E). These findings are consistent with the metastatic progression in a clinical setting, as liver metastasis is associated with worse outcomes and neuroendocrine differentiation in patients ^49^. Brain metastases, while rare in PCa overall, may be more common in NEPC due to its neuroendocrine and small cell-like features ^50^. Immunohistochemistry was performed using a chromogranin antibody on brain and liver sections derived from placebo and GSNO-treated mice. Results showed a significant reduction in chromogranin expression in GSNO-treated brains and liver (Figure 6F), supporting the observations made in Figure 6D. Next, western blot and immunohistochemistry were performed on the sections derived from tumor grafts of placebo controls and GSNO-treated mice. Results revealed a significant reduction in the expression of neuroendocrine markers MYCN, cMYC, Synaptophysin, Chromogranin, and ER stress markers such as PERK, BIP, pIRE1 alpha, PDI, and CHOP, respectively, in the GSNO-treated samples (Figure 6G-H). These observations highlight the role of ER stress in NEPC transdifferentiation and, therefore, in delineating the metastatic landscape. To further evaluate the prognostic significance of ER Stress Markers in PCa metastasis, we performed Gene Expression Profiling Interactive Analysis (GEPIA). GEPIA showed that high expression of several ER stress markers Is associated with an increased risk of distant metastasis in men with PCa: MYCN: Hazard ratio (HR) for relapse = 2.1, p = 0.0018; DDIT3: HR for relapse = 2.3, p = 0.00019; PDI (Protein Disulfide Isomerase: HR for relapse = 2.0, p = 0.0012 and NOS3: HR for relapse = 1.7, p = 0.015 respectively. NOS3 involvement suggests a potential link between NO signaling dysregulation and metastasis(Supplementary Figure 7). To confirm these observations, we evaluated the expression of markers of epithelial to mesenchymal transition in the protein lysates derived from murine tumor grafts from Figure 6A. Results showed that NO supplementation, which was found to inhibit ER stress response markers, was able to significantly reduce the expression of markers such as TGFb, SNAI1, SNAI2, NCADH, ZEB1, Claudin, and ZO1, respectively (Figure 6I). Together, these results indicate that higher expression of ER stress markers is associated with a greater risk of distant metastasis in high-grade PCa patients and NO supplementation can inhibit it.

### NO induces S-nitrosylation of MYCN to inhibit ER stress responses

One of the primary mechanisms by which NO exerts its cellular effects is through S-nitrosylation, a process wherein a nitroso moiety from NO-derived metabolites covalently attaches to a reactive cysteine residue, forming an S-nitrosothiol (SNO) ^51,52^. Given the significant inhibitory effects of NO supplementation on MYCN-induced ER stress, we hypothesized that S-nitrosylation of MYCN by GSNO (S-Nitrosoglutathione) is behind dampening the ER stress response. Using the GPS-SNO 1.0 software, we identified 16 cysteine residues on the MYCN protein, conforming to the acid-base nitrosylation conservative motif. Among these, Cys4, Cys186, and Cys464 showed the highest predicted thresholds for S-nitrosylation (cutoff 2.443) (Figure 7A and Supplementary Figure 8A). To confirm that GSNO induces S-nitrosylation of MYCN, we employed a biotin-switch assay on protein lysates derived from tumor samples of mice treated with or without GSNO, which demonstrated the formation of S-nitrosylated MYCN upon GSNO treatment, specifically in the heavy chain of MYCN (Figure 7B). This highlights that NO supplementation can modify MYCN through S-nitrosylation, altering its function.

**FIGURE 7.**
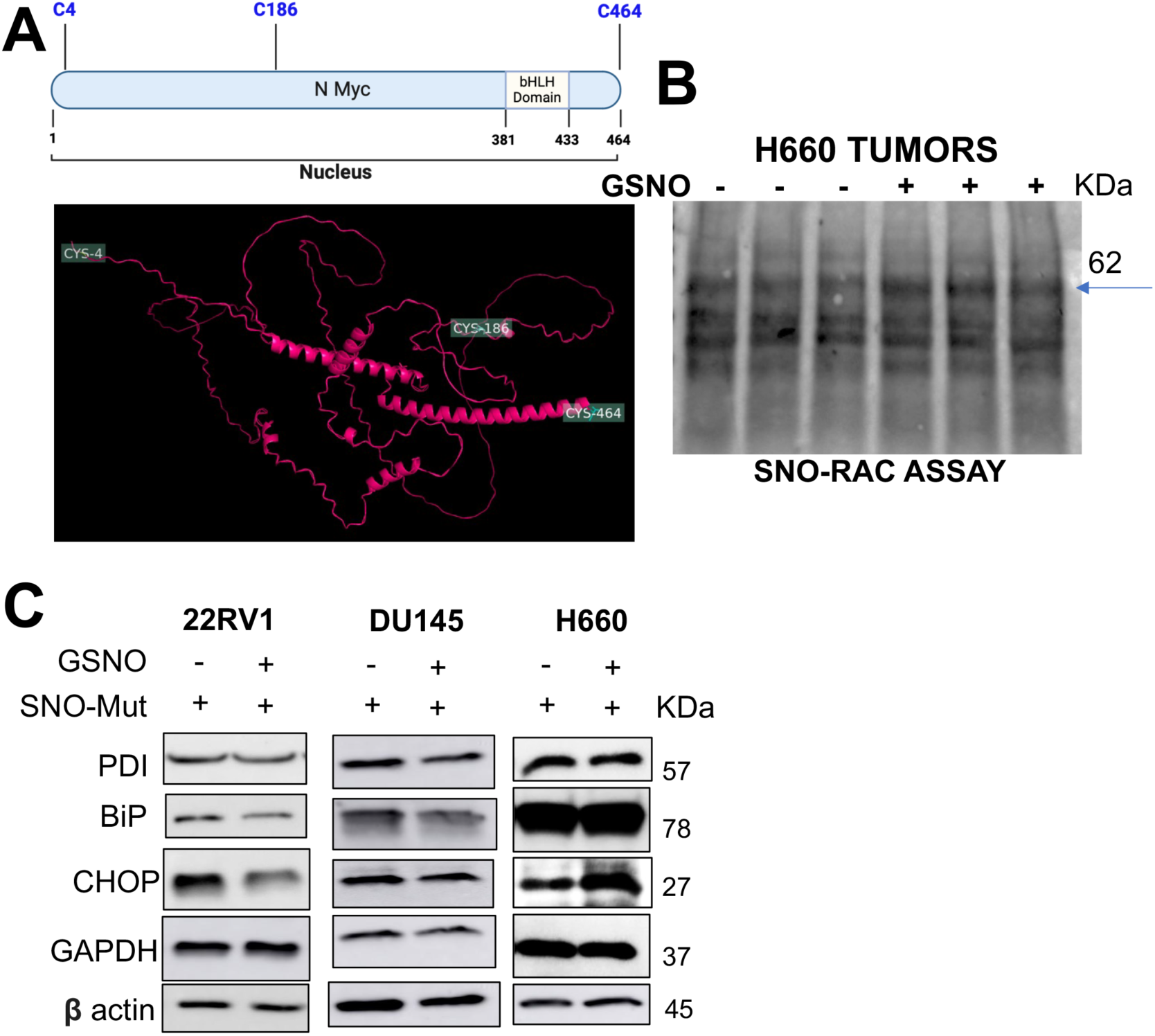
S-nitrosylation of MYCN by GSNO and its effects on protein binding and ER stress in NEPC cells. **(a)** GPS-SNO 1.0 software predicted 16 cysteine residues on the MYCN protein susceptible to S-nitrosylation, with Cys4, Cys186, and Cys464 exhibiting the highest thresholds for nitrosylation (cutoff 2.443). **(b)** Biotin-switch assay confirmed S-nitrosylation of MYCN upon GSNO treatment, specifically affecting the heavy chain of MYCN. **(c)** Western blot showing the protein expression of ER stress markers (PDI, BIP, and CHOP) in 22Rv1, DU145, and H660 cells, post-transfection with MYCN plasmid-containing mutations at Cys4, Cys186, and Cys464 sites, followed by GSNO treatment (50ug) for 48 hours.

To investigate whether S-nitrosylation of MYCN at specific cysteine residues contributes to NO-mediated suppression of ER stress in NEPC, we generated site-directed mutations at Cys4, Cys186, and Cys464 (Supplementary Figure 8B). Structural analysis of MYCN revealed the topological locations of these cysteines (Figure 7A), which may influence MYCN’s functional domains. We introduced these mutated MYCN constructs into 22RV1, DU145, and H660 cell lines. Following transfection, cells were exposed to vehicle or GSNO (50 μM) treatment for 48 hours, after which protein lysates were analyzed via western blot for the ER stress markers. The results (Figure 7C) demonstrated that mutations at S-nitrosylation sites on MYCN prevented NO-induced ER stress inhibition in H660 cells, whereas no such effect was observed in 22RV1 or DU145 cells. These findings strongly suggest that NO specifically targets MYCN to suppress ER stress responses in NEPC, with S-nitrosylation at Cys4, Cys186, and Cys464 playing a crucial role in this selective regulatory mechanism.

We further hypothesized that NO-induced S-nitrosylation could interfere with the binding of various protein partners to MYCN, thus impairing its ability to promote tumor progression in NEPC cells. To explore this, we conducted proteomic analysis through IP-MS in GSNO-treated NEPC xenografts using antibody against MYCN (Figure 8A). Analysis outcomes showed a substantial reduction in MYCN-binding proteins linked to RNA and DNA metabolism, ER protein import, CFTR misfolding and degradation, transcriptional regulation, and chromatin remodeling (Figure 8B). Figure 8C-F details the spectrum of inhibited interactions, with notable declines in co-translational ER import processes and protein-folding functions. This pattern underscores a targeted disruption in MYCN’s capacity to maintain proteostasis within the ER, which is further corroborated by the decreased presence of protein partners involved in misfolding management and chromatin structure regulation. The data suggest that NO-induced S-nitrosylation of MYCN reconfigures the ER’s adaptive response, enhancing protein-folding efficiency while limiting the accumulation of misfolded proteins that typically contribute to stress. These modifications to ER function are clinically relevant, as they point to a reduction in the UPR threshold in NEPC cells, thereby lessening the cells’ ability to counterbalance stress-induced damage and maintain lineage plasticity. By compromising MYCN’s control over these cellular networks, NO treatment disrupts NEPC cells’ resilience and adaptability—a key factor in their survival and resistance to standard treatments. These findings highlight the therapeutic potential of targeting MYCN S-nitrosylation to destabilize the cellular equilibrium essential for NEPC progression, offering a strategic avenue for reducing therapeutic resistance in this aggressive cancer phenotype.

**Figure 8:**
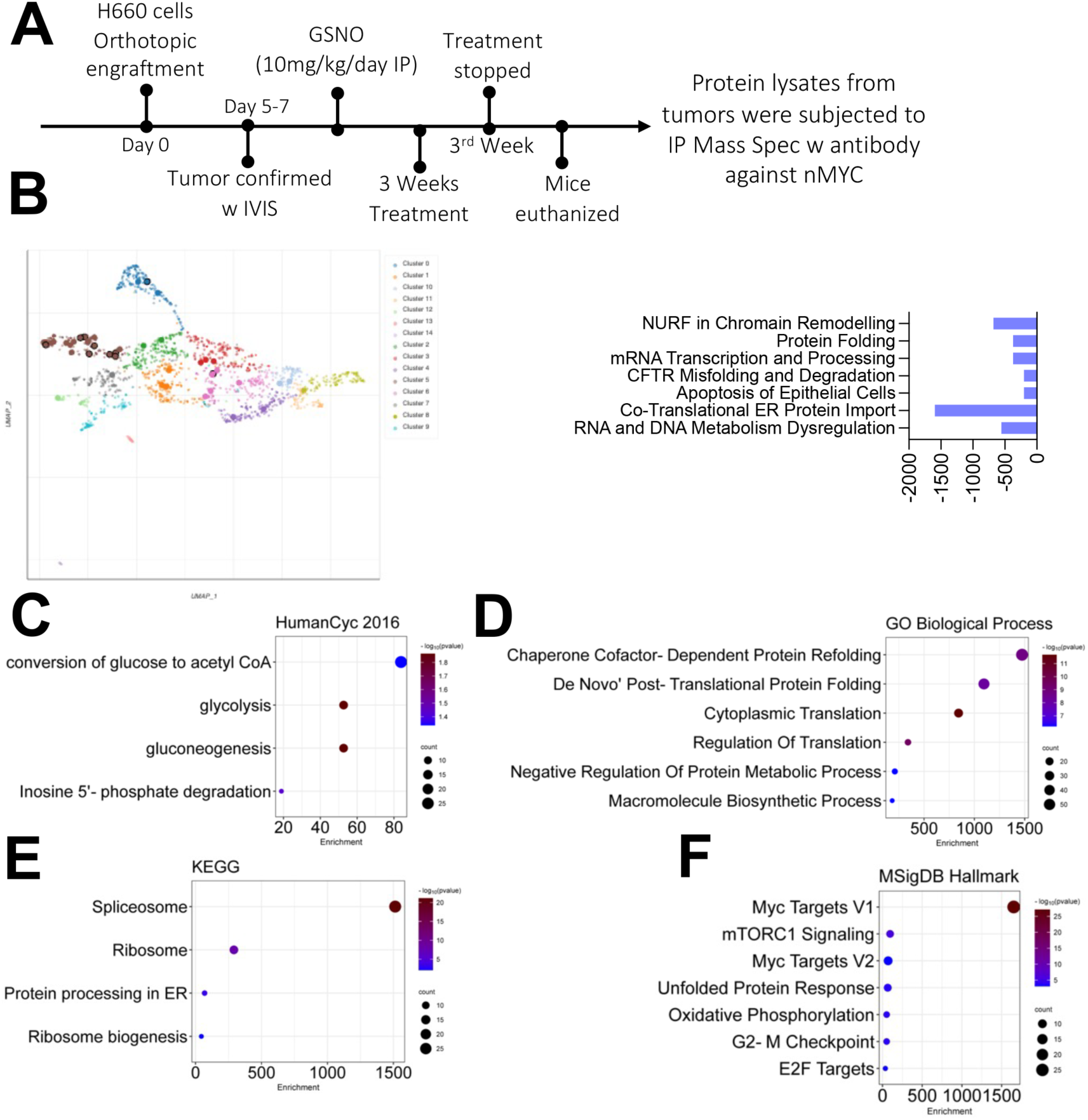
Proteomic Analysis of GSNO Treatment Effects on Protein Interactions in NEPC tumors: **(a)** Schematic illustration of the immunoprecipitation followed by mass spectrometry (IP-MS) protocol used to analyze protein interactions in control and GSNO-treated mice. **(b)** Comparison of protein interactions between control and GSNO-treated mice. NO supplementation via GSNO inhibited the binding of proteins involved in key cellular processes such as RNA/DNA metabolism, ER protein import, Protein folding, and Chromatin remodeling. (c-f) Gene Ontology and pathway analyses were performed on down-regulated proteins in nuclear lysates from tumor and GSNO-treated tumor samples. The results are presented in bubble plots, where bubble size represents the number of genes and color intensity indicates statistical significance. **(c)** Human Cyt 2016: Bubble plot illustrating enriched cytological terms associated with down-regulated proteins. This analysis provides insights into the cellular compartments and structures affected by GSNO treatment. **(d)** GO Biological Process: Bubble plot showing enriched cellular processes impacted by GSNO treatment. This analysis highlights the biological functions most significantly altered in response to NO supplementation. **(e)** KEGG Pathway Analysis: Bubble plot depicting enriched cellular pathways affected by GSNO treatment. This analysis reveals the broader cellular systems and networks influenced by NO-mediated protein interaction changes. **(f)** MSigDB Hallmark: Bubble plot showing enrichment of hallmark gene sets, providing a high-level overview of the cellular states and processes altered by GSNO treatment

## DISCUSSION

NEPC remains a formidable challenge in the clinical management of PCa due to its aggressive nature and resistance to conventional therapies^1,2,5,6^. ER stress is a pivotal factor in NEPC progression, as it facilitates tumor cell survival under hostile metabolic conditions. MYCN is a well-known oncogenic driver of neuroendocrine differentiation ^3,4,53–63^. However, its role in contributing to the increased biosynthetic demands of neuroendocrine cancer cells remains understudied. This information is essential because sustained ER stress leads to increased Ca^2+^ release, mitochondrial dysfunction, and the generation of ROS, which in turn exacerbates ER stress, creating a vicious cycle that promotes tumor growth and metastasis ^9–12^. Understanding how MYCN, can play an essential role in delineating ER stress responses and other metabolic changes within an NEPC cell could assist in effectively targeting NEPC cancers. Our study sheds light on the molecular underpinnings of MYCN-driven ER stress and demonstrates how the dysregulation of a protein modification, S-nitrosylation allows the MYCN to increase the ER stress in an NEPC cell. We demonstrated that overcoming this limitation through exogenous supplementation of nitric oxide, a molecule whose production otherwise is reduced in high-grade PCa, can reinstate the S-nitrosylation of MYCN at specific sites (Cys4, Cys186, Cys46). The positive effects of NO treatment are observed not only in reducing the tumor burden of NEPC in murine models but also in reducing the overall metastasis to the Brain and Liver. This observation aligns with the findings of retrospective studies, which state that NEPC patients have significantly more frequent visceral metastases compared to patients with castration-resistant adenocarcinoma (62% vs 24%, p < 0.001) with Liver and Brain metastases, particularly common in NEPC ^1,64^. Furthermore, our *in vitro* studies elucidated how NO supplementation can reverse the effects of ER stress by the suppression of key unfolded protein response (UPR) markers, a survival mechanism to cope with increased protein folding demands and cellular stress; reducing the release of Ca^2+^ from ER to the cytoplasm, thereby inhibiting the overall glycolytic stress in the NEPC cells; ultimately inhibiting the colony forming and proliferative properties of NEPC cells.

The therapeutic implications of these findings are significant. Given the poor response of NEPC to standard chemotherapies and emerging chemo-immunotherapy combinations^1,2,5,6^, our data highlight NO supplementation as a promising therapeutic strategy. By targeting the nitroso-redox imbalance and restoring S-nitrosylation in NEPC cells, NO donors like GSNO could potentially overcome one of the key mechanisms of therapy resistance—namely, the unchecked activity of MYCN. This approach not only addresses the metabolic and biosynthetic imbalances driven by MYCN overexpression but also restores mitochondrial function and inhibits the ER stress that underpins NEPC aggressiveness. Moreover, the role of NO in inhibiting lineage plasticity offers a novel approach to halting the transition of PCa cells to a neuroendocrine phenotype. As lineage plasticity remains one of the most intractable problems in NEPC, disrupting this process through the modulation of MYCN function could pave the way for more effective treatment regimens.

While the results of our study provide robust evidence for the therapeutic potential of NO-based therapies in NEPC, there are several limitations to consider. First, the extent to which NO supplementation can be integrated into existing therapeutic regimens needs to be further explored. Combination therapies involving NO donors and current chemo-immunotherapies may offer synergistic benefits, but this hypothesis remains to be tested in clinical trials. Additionally, while our *in-vivo* models showed promising reductions in tumor burden, the long-term effects of NO supplementation on NEPC progression, especially regarding potential resistance mechanisms, require further investigation.

Future research should focus on understanding the broader implications of S-nitrosylation in regulating other oncogenic drivers beyond MYCN. It is also essential to investigate the role of NO in modulating the tumor microenvironment (TME), particularly in terms of immune cell infiltration and the potential reprogramming of tumor-associated macrophages (TAMs), which could further enhance the anti-tumor efficacy of NO-based therapies.

In conclusion, our study uncovers a novel mechanism by which ER stress and MYCN-mediated lineage plasticity drive NEPC progression. The therapeutic potential of NO supplementation to restore S-nitrosylation, re-regulate ER stress responses, and inhibit the oncogenic activity of MYCN presents a promising avenue for addressing the treatment resistance and aggressiveness of NEPC. This approach may offer a new strategy for improving outcomes in patients with this challenging form of prostate cancer.

## Resource availability

### Lead contact

All data, codes, and materials used in the study are available to any researcher for reproducing or extending the analysis. Materials transfer agreements (MTAs) will be required to request access. RNA sequencing data have been deposited in a public database.

Further requests for information should be directed to and will be fulfilled by the lead contact, Himanshu Arora.

## Acknowledgments

We thank all the peers (Drs. Dean Tang, Robert Matusik) and collaborators (Drs. Bonnie Bloomberg, Zhibin Chen, Fangliang Zhang) for their insights, suggestions, and support during this study.

## Author Contributions

HA conceived the project, and HA and RQ designed the experiments. HA, FF, OCN, RQ, SK, and RAD performed the experiments with the assistance of AE, KS, DL, and RG. HA, RQ, FF, and OCN analyzed and interpreted the data. DVB and SS performed the bioinformatic analysis. HA and RQ wrote the manuscript. HA, JMH and DL involved in funding acquisition. All authors had the opportunity to discuss the results and comment on the manuscript.

## STAR+METHODS

Detailed methods are provided in the online version of this paper and include the following:

KEY RESOURCES TABLE

METHOD DETAILS STATISTICAL ANALYSIS

## References

1. Yamada, Y., and Beltran, H. (2021). Clinical and Biological Features of Neuroendocrine Prostate Cancer. Curr Oncol Rep 23, 15. 10.1007/s11912-020-01003-9.

2. Alabi, B.R., Liu, S., and Stoyanova, T. (2022). Current and emerging therapies for neuroendocrine prostate cancer. Pharmacol Ther 238, 108255. 10.1016/j.pharmthera.2022.108255.

3. Zamora, I., Freeman, M.R., Encío, I.J., and Rotinen, M. (2023). Targeting Key Players of Neuroendocrine Differentiation in Prostate Cancer. Int J Mol Sci 24. 10.3390/ijms241813673.

4. Okasho, K., Ogawa, O., and Akamatsu, S. (2021). Narrative review of challenges in the management of advanced neuroendocrine prostate cancer. Transl Androl Urol 10, 3953–3962. 10.21037/tau-20-1131.

5. Alonso-Gordoa, T., Goodman, M., Vulsteke, C., Roubaud, G., Zhang, J., Parikh, M., Piulats, J.M., Azaro, A., James, G.D., Cavazzina, R., et al. (2024). A phase II study (AARDVARC) of AZD4635 in combination with durvalumab and cabazitaxel in patients with progressive, metastatic, castration-resistant prostate cancer. ESMO Open 9, 103446. 10.1016/j.esmoop.2024.103446.

6. Brown, L.C., Halabi, S., Somarelli, J.A., Humeniuk, M., Wu, Y., Oyekunle, T., Howard, L., Huang, J., Anand, M., Davies, C., et al. (2022). A phase 2 trial of avelumab in men with aggressive-variant or neuroendocrine prostate cancer. Prostate Cancer and Prostatic Diseases 25, 762–769. 10.1038/s41391-022-00524-7.

7. Bravo, R., Parra, V., Gatica, D., Rodriguez, A.E., Torrealba, N., Paredes, F., Wang, Z.V., Zorzano, A., Hill, J.A., Jaimovich, E., et al. (2013). Endoplasmic reticulum and the unfolded protein response: dynamics and metabolic integration. Int Rev Cell Mol Biol 301, 215–290. 10.1016/b978-0-12-407704-1.00005-1.

8. Chen, X., and Cubillos-Ruiz, J.R. (2021). Endoplasmic reticulum stress signals in the tumour and its microenvironment. Nat Rev Cancer 21, 71–88. 10.1038/s41568-020-00312-2.

9. Cao, S.S., and Kaufman, R.J. (2014). Endoplasmic reticulum stress and oxidative stress in cell fate decision and human disease. Antioxid Redox Signal 21, 396–413. 10.1089/ars.2014.5851.

10. Bhattarai, K.R., Riaz, T.A., Kim, H.R., and Chae, H.J. (2021). The aftermath of the interplay between the endoplasmic reticulum stress response and redox signaling. Exp Mol Med 53, 151–167. 10.1038/s12276-021-00560-8.

11. Chen, X., Shi, C., He, M., Xiong, S., and Xia, X. (2023). Endoplasmic reticulum stress: molecular mechanism and therapeutic targets. Signal Transduction and Targeted Therapy 8, 352. 10.1038/s41392-023-01570-w.

12. Morris, G., Puri, B.K., Walder, K., Berk, M., Stubbs, B., Maes, M., and Carvalho, A.F. (2018). The Endoplasmic Reticulum Stress Response in Neuroprogressive Diseases: Emerging Pathophysiological Role and Translational Implications. Molecular Neurobiology 55, 8765–8787. 10.1007/s12035-018-1028-6.

13. Berger, A., Brady, N.J., Bareja, R., Robinson, B., Conteduca, V., Augello, M.A., Puca, L., Ahmed, A., Dardenne, E., Lu, X., et al. (2019). N-Myc-mediated epigenetic reprogramming drives lineage plasticity in advanced prostate cancer. J Clin Invest 129, 3924–3940. 10.1172/JCI127961.

14. Davies, A., Zoubeidi, A., and Selth, L.A. (2020). The epigenetic and transcriptional landscape of neuroendocrine prostate cancer. Endocr Relat Cancer 27, R35–R50. 10.1530/ERC-19-0420.

15. Qiu, X., Boufaied, N., Hallal, T., Feit, A., de Polo, A., Luoma, A.M., Alahmadi, W., Larocque, J., Zadra, G., Xie, Y., et al. (2022). MYC drives aggressive prostate cancer by disrupting transcriptional pause release at androgen receptor targets. Nat Commun 13, 2559. 10.1038/s41467-022-30257-z.

16. Yasumizu, Y., Rajabi, H., Jin, C., Hata, T., Pitroda, S., Long, M.D., Hagiwara, M., Li, W., Hu, Q., Liu, S., et al. (2020). MUC1-C regulates lineage plasticity driving progression to neuroendocrine prostate cancer. Nat Commun 11, 338. 10.1038/s41467-019-14219-6.

17. Lee, J.K., Phillips, J.W., Smith, B.A., Park, J.W., Stoyanova, T., McCaffrey, E.F., Baertsch, R., Sokolov, A., Meyerowitz, J.G., Mathis, C., et al. (2016). N-Myc Drives Neuroendocrine Prostate Cancer Initiated from Human Prostate Epithelial Cells. Cancer Cell 29, 536–547. 10.1016/j.ccell.2016.03.001.

18. 18. Firdaus, F., Kuchakulla, M., Qureshi, R., Dulce, R.A., Soni, Y., Van Booven, D.J., Shah, K., Masterson, T., Rosete, O.J., Punnen, S., et al. (2022). S-nitrosylation of CSF1 receptor increases the efficacy of CSF1R blockage against prostate cancer. Cell Death Dis 13, 859. 10.1038/s41419-022-05289-4.

19. Storck, W.K., May, A.M., Westbrook, T.C., Duan, Z., Morrissey, C., Yates, J.A., and Alumkal, J.J. (2022). The Role of Epigenetic Change in Therapy-Induced Neuroendocrine Prostate Cancer Lineage Plasticity. Front Endocrinol (Lausanne) 13, 926585. 10.3389/fendo.2022.926585.

20. Wishahi, M. (2024). Treatment-induced neuroendocrine prostate cancer and de novo neuroendocrine prostate cancer: Identification, prognosis and survival, genetic and epigenetic factors. World J Clin Cases 12, 2143–2146. 10.12998/wjcc.v12.i13.2143.

21. Zhou, Z., Wang, Q., and Michalak, M. (2021). Inositol Requiring Enzyme (IRE), a multiplayer in sensing endoplasmic reticulum stress. Anim Cells Syst (Seoul) 25, 347–357. 10.1080/19768354.2021.2020901.

22. Kepp, O., Bezu, L., and Kroemer, G. (2022). The endoplasmic reticulum chaperone BiP: a target for immunogenic cell death inducers? Oncoimmunology 11, 2092328. 10.1080/2162402X.2022.2092328.

23. Robin, M.J.D., Appelman, M.D., Vos, H.R., van Es, R.M., Paton, J.C., Paton, A.W., Burgering, B., Fickert, P., Heijmans, J., and van de Graaf, S.F.J. (2018). Calnexin Depletion by Endoplasmic Reticulum Stress During Cholestasis Inhibits the Na(+)-Taurocholate Cotransporting Polypeptide. Hepatol Commun 2, 1550–1566. 10.1002/hep4.1262.

24. Hu, H., Tian, M., Ding, C., and Yu, S. (2018). The C/EBP Homologous Protein (CHOP) Transcription Factor Functions in Endoplasmic Reticulum Stress-Induced Apoptosis and Microbial Infection. Front Immunol 9, 3083. 10.3389/fimmu.2018.03083.

25. Varone, E., Decio, A., Chernorudskiy, A., Minoli, L., Brunelli, L., Ioli, F., Piotti, A., Pastorelli, R., Fratelli, M., Gobbi, M., et al. (2021). The ER stress response mediator ERO1 triggers cancer metastasis by favoring the angiogenic switch in hypoxic conditions. Oncogene 40, 1721–1736. 10.1038/s41388-021-01659-y.

26. Chen, X., Shi, C., He, M., Xiong, S., and Xia, X. (2023). Endoplasmic reticulum stress: molecular mechanism and therapeutic targets. Signal Transduct Target Ther 8, 352. 10.1038/s41392-023-01570-w.

27. Wu, M.J., Chen, C.J., Lin, T.Y., Liu, Y.Y., Tseng, L.L., Cheng, M.L., Chuu, C.P., Tsai, H.K., Kuo, W.L., Kung, H.J., and Wang, W.C. (2021). Targeting KDM4B that coactivates c-Myc-regulated metabolism to suppress tumor growth in castration-resistant prostate cancer. Theranostics 11, 7779–7796. 10.7150/thno.58729.

28. Gao, L., Schwartzman, J., Gibbs, A., Lisac, R., Kleinschmidt, R., Wilmot, B., Bottomly, D., Coleman, I., Nelson, P., McWeeney, S., and Alumkal, J. (2013). Androgen receptor promotes ligand-independent prostate cancer progression through c-Myc upregulation. PLoS One 8, e63563. 10.1371/journal.pone.0063563.

29. Rabender, C.S., Alam, A., Sundaresan, G., Cardnell, R.J., Yakovlev, V.A., Mukhopadhyay, N.D., Graves, P., Zweit, J., and Mikkelsen, R.B. (2015). The Role of Nitric Oxide Synthase Uncoupling in Tumor Progression. Mol Cancer Res 13, 1034–1043. 10.1158/1541-7786.MCR-15-0057-T.

30. Goncalves, D.A., Jasiulionis, M.G., and Melo, F.H.M. (2021). The Role of the BH4 Cofactor in Nitric Oxide Synthase Activity and Cancer Progression: Two Sides of the Same Coin. Int J Mol Sci 22. 10.3390/ijms22179546.

31. Barnett, S.D., and Buxton, I.L.O. (2017). The role of S-nitrosoglutathione reductase (GSNOR) in human disease and therapy. Crit Rev Biochem Mol Biol 52, 340–354. 10.1080/10409238.2017.1304353.

32. Werner, E.R., Gorren, A.C., Heller, R., Werner-Felmayer, G., and Mayer, B. (2003). Tetrahydrobiopterin and nitric oxide: mechanistic and pharmacological aspects. Exp Biol Med (Maywood) 228, 1291–1302. 10.1177/153537020322801108.

33. Berka, V., Yeh, H.C., Gao, D., Kiran, F., and Tsai, A.L. (2004). Redox function of tetrahydrobiopterin and effect of L-arginine on oxygen binding in endothelial nitric oxide synthase. Biochemistry 43, 13137–13148. 10.1021/bi049026j.

34. Groenendyk, J., Agellon, L.B., and Michalak, M. (2021). Calcium signaling and endoplasmic reticulum stress. Int Rev Cell Mol Biol 363, 1–20. 10.1016/bs.ircmb.2021.03.003.

35. Bery, F., Cancel, M., Chantome, A., Guibon, R., Bruyere, F., Rozet, F., Maheo, K., and Fromont, G. (2020). The Calcium-Sensing Receptor is A Marker and Potential Driver of Neuroendocrine Differentiation in Prostate Cancer. Cancers (Basel) 12. 10.3390/cancers12040860.

36. Matuz-Mares, D., Gonzalez-Andrade, M., Araiza-Villanueva, M.G., Vilchis-Landeros, M.M., and Vazquez-Meza, H. (2022). Mitochondrial Calcium: Effects of Its Imbalance in Disease. Antioxidants (Basel) 11. 10.3390/antiox11050801.

37. Wang, Y., Wang, Y., Ci, X., Choi, S.Y.C., Crea, F., Lin, D., and Wang, Y. (2021). Molecular events in neuroendocrine prostate cancer development. Nat Rev Urol 18, 581–596. 10.1038/s41585-021-00490-0.

38. Yin, Y., Xu, L., Chang, Y., Zeng, T., Chen, X., Wang, A., Groth, J., Foo, W.C., Liang, C., Hu, H., and Huang, J. (2019). N-Myc promotes therapeutic resistance development of neuroendocrine prostate cancer by differentially regulating miR-421/ATM pathway. Mol Cancer 18, 11. 10.1186/s12943-019-0941-2.

39. Smith, M.I., and Deshmukh, M. (2007). Endoplasmic reticulum stress-induced apoptosis requires bax for commitment and Apaf-1 for execution in primary neurons. Cell Death Differ 14, 1011–1019. 10.1038/sj.cdd.4402089.

40. Dufey, E., Bravo-San Pedro, J.M., Eggers, C., Gonzalez-Quiroz, M., Urra, H., Sagredo, A.I., Sepulveda, D., Pihan, P., Carreras-Sureda, A., Hazari, Y., et al. (2020). Genotoxic stress triggers the activation of IRE1alpha-dependent RNA decay to modulate the DNA damage response. Nat Commun 11, 2401. 10.1038/s41467-020-15694-y.

41. Wang, P., Liao, H., Wang, Q., Xie, H., Wang, H., Yang, M., and Liu, S. (2022). L1 Syndrome Prenatal Diagnosis Supplemented by Functional Analysis of One L1CAM Gene Missense Variant. Reprod Sci 29, 768–780. 10.1007/s43032-021-00828-4.

42. Liu, Z., Li, C., Kang, N., Malhi, H., Shah, V.H., and Maiers, J.L. (2019). Transforming growth factor beta (TGFbeta) cross-talk with the unfolded protein response is critical for hepatic stellate cell activation. J Biol Chem 294, 3137–3151. 10.1074/jbc.RA118.005761.

43. Jaud, M., Philippe, C., Di Bella, D., Tang, W., Pyronnet, S., Laurell, H., Mazzolini, L., Rouault-Pierre, K., and Touriol, C. (2020). Translational Regulations in Response to Endoplasmic Reticulum Stress in Cancers. Cells 9. 10.3390/cells9030540.

44. Guan, B.J., Krokowski, D., Majumder, M., Schmotzer, C.L., Kimball, S.R., Merrick, W.C., Koromilas, A.E., and Hatzoglou, M. (2014). Translational control during endoplasmic reticulum stress beyond phosphorylation of the translation initiation factor eIF2alpha. J Biol Chem 289, 12593–12611. 10.1074/jbc.M113.543215.

45. Shaheen, A. (2018). Effect of the unfolded protein response on ER protein export: a potential new mechanism to relieve ER stress. Cell Stress Chaperones 23, 797–806. 10.1007/s12192-018-0905-2.

46. Han, J., and Kaufman, R.J. (2016). The role of ER stress in lipid metabolism and lipotoxicity. J Lipid Res 57, 1329–1338. 10.1194/jlr.R067595.

47. Beltran, H., Rickman, D.S., Park, K., Chae, S.S., Sboner, A., MacDonald, T.Y., Wang, Y., Sheikh, K.L., Terry, S., Tagawa, S.T., et al. (2011). Molecular characterization of neuroendocrine prostate cancer and identification of new drug targets. Cancer Discov 1, 487–495. 10.1158/2159-8290.CD-11-0130.

48. Shahryari, V., Nip, H., Saini, S., Dar, A.A., Yamamura, S., Mitsui, Y., Colden, M., Bucay, N., Tabatabai, L.Z., Greene, K., et al. (2016). Pre-clinical Orthotopic Murine Model of Human Prostate Cancer. J Vis Exp. 10.3791/54125.

49. Cloyd, J.M., Ejaz, A., Konda, B., Makary, M.S., and Pawlik, T.M. (2020). Neuroendocrine liver metastases: a contemporary review of treatment strategies. Hepatobiliary Surg Nutr 9, 440–451. 10.21037/hbsn.2020.04.02.

50. Krug, S., Teupe, F., Michl, P., Gress, T.M., and Rinke, A. (2019). Brain metastases in patients with neuroendocrine neoplasms: risk factors and outcome. BMC Cancer 19, 362. 10.1186/s12885-019-5559-7.

51. Nelson, E.J., Connolly, J., and McArthur, P. (2003). Nitric oxide and S-nitrosylation: excitotoxic and cell signaling mechanism. Biol Cell 95, 3–8. 10.1016/s0248-4900(03)00004-2.

52. Beigi, F., Gonzalez, D.R., Minhas, K.M., Sun, Q.A., Foster, M.W., Khan, S.A., Treuer, A.V., Dulce, R.A., Harrison, R.W., Saraiva, R.M., et al. (2012). Dynamic denitrosylation via S-nitrosoglutathione reductase regulates cardiovascular function. Proc Natl Acad Sci U S A 109, 4314–4319. 10.1073/pnas.1113319109.

53. Braglia, L., Zavatti, M., Vinceti, M., Martelli, A.M., and Marmiroli, S. (2020). Deregulated PTEN/PI3K/AKT/mTOR signaling in prostate cancer: Still a potential druggable target? Biochim Biophys Acta Mol Cell Res 1867, 118731. 10.1016/j.bbamcr.2020.118731.

54. Damodaran, S., Damaschke, N., Gawdzik, J., Yang, B., Shi, C., Allen, G.O., Huang, W., Denu, J., and Jarrard, D. (2017). Dysregulation of Sirtuin 2 (SIRT2) and histone H3K18 acetylation pathways associates with adverse prostate cancer outcomes. BMC Cancer 17, 874. 10.1186/s12885-017-3853-9.

55. Faugeroux, V., Pailler, E., Oulhen, M., Deas, O., Brulle-Soumare, L., Hervieu, C., Marty, V., Alexandrova, K., Andree, K.C., Stoecklein, N.H., et al. (2020). Genetic characterization of a unique neuroendocrine transdifferentiation prostate circulating tumor cell-derived eXplant model. Nat Commun 11, 1884. 10.1038/s41467-020-15426-2.

56. Lee, J.H., Yang, B., Lindahl, A.J., Damaschke, N., Boersma, M.D., Huang, W., Corey, E., Jarrard, D.F., and Denu, J.M. (2017). Identifying Dysregulated Epigenetic Enzyme Activity in Castrate-Resistant Prostate Cancer Development. ACS Chem Biol 12, 2804–2814. 10.1021/acschembio.6b01035.

57. Ribarska, T., Bastian, K.M., Koch, A., and Schulz, W.A. (2012). Specific changes in the expression of imprinted genes in prostate cancer--implications for cancer progression and epigenetic regulation. Asian J Androl 14, 436–450. 10.1038/aja.2011.160.

58. Schagdarsurengin, U., Lammert, A., Schunk, N., Sheridan, D., Gattenloehner, S., Steger, K., Wagenlehner, F., and Dansranjavin, T. (2017). Impairment of IGF2 gene expression in prostate cancer is triggered by epigenetic dysregulation of IGF2-DMR0 and its interaction with KLF4. Cell Commun Signal 15, 40. 10.1186/s12964-017-0197-7.

59. Yu, T., Wang, C., Yang, J., Guo, Y., Wu, Y., and Li, X. (2017). Metformin inhibits SUV39H1-mediated migration of prostate cancer cells. Oncogenesis 6, e324. 10.1038/oncsis.2017.28.

60. Kaarijärvi, R., Kaljunen, H., and Ketola, K. (2021). Molecular and Functional Links between Neurodevelopmental Processes and Treatment-Induced Neuroendocrine Plasticity in Prostate Cancer Progression. Cancers (Basel) 13. 10.3390/cancers13040692.

61. Imamura, J., Ganguly, S., Muskara, A., Liao, R.S., Nguyen, J.K., Weight, C., Wee, C.E., Gupta, S., and Mian, O.Y. (2023). Lineage plasticity and treatment resistance in prostate cancer: the intersection of genetics, epigenetics, and evolution. Front Endocrinol (Lausanne) 14, 1191311. 10.3389/fendo.2023.1191311.

62. Wei, G., Zhang, X., Liu, S., Hou, W., and Dai, Z. (2024). Comprehensive data mining reveals RTK/RAS signaling pathway as a promoter of prostate cancer lineage plasticity through transcription factors and CNV. Scientific Reports 14, 11688. 10.1038/s41598-024-62256-z.

63. Beltran, H., Hruszkewycz, A., Scher, H.I., Hildesheim, J., Isaacs, J., Yu, E.Y., Kelly, K., Lin, D., Dicker, A., Arnold, J., et al. (2019). The Role of Lineage Plasticity in Prostate Cancer Therapy Resistance. Clin Cancer Res 25, 6916–6924. 10.1158/1078-0432.Ccr-19-1423.

64. Suzuki, K., Terakawa, T., Jimbo, N., Inaba, R., Nakano, Y., and Fujisawa, M. (2020). Clinical Features of Treatment-related Neuroendocrine Prostate Cancer: A Case Series. Anticancer Res 40, 3519–3526. 10.21873/anticanres.14340.

