## Supplementary material for "Targeting Endoplasmic Reticulum Stress and Nitroso-Redox Imbalance in Neuroendocrine Prostate Cancer: The Therapeutic Role of Nitric Oxide": Supp Material

**Supp. Figure 1.** The expression of ER stress markers in PCa patients' data was obtained from the Cancer genome atlas data (TCGA).

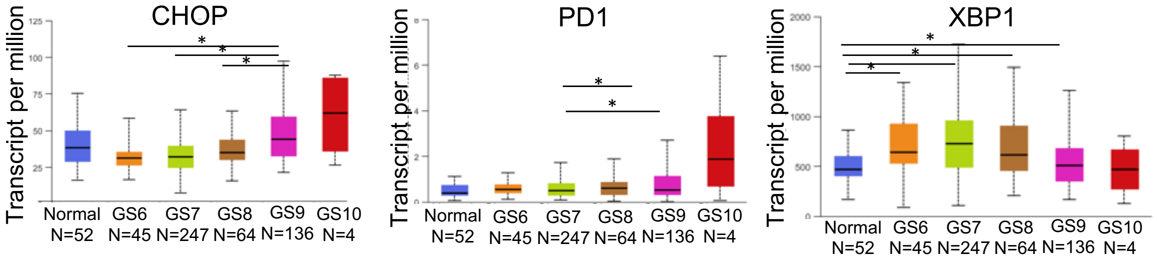

**Supp. Figure 2.** (a) Experimental steps are taken to generate MyCaP, MyCaPshAR, and MyCaPAPIPC cells, respectively. (b) shows expression of AR and MYCN in MyCaP, MyCaP$^{shAR}$ and MyCaP$^{APIPC}$ cells.

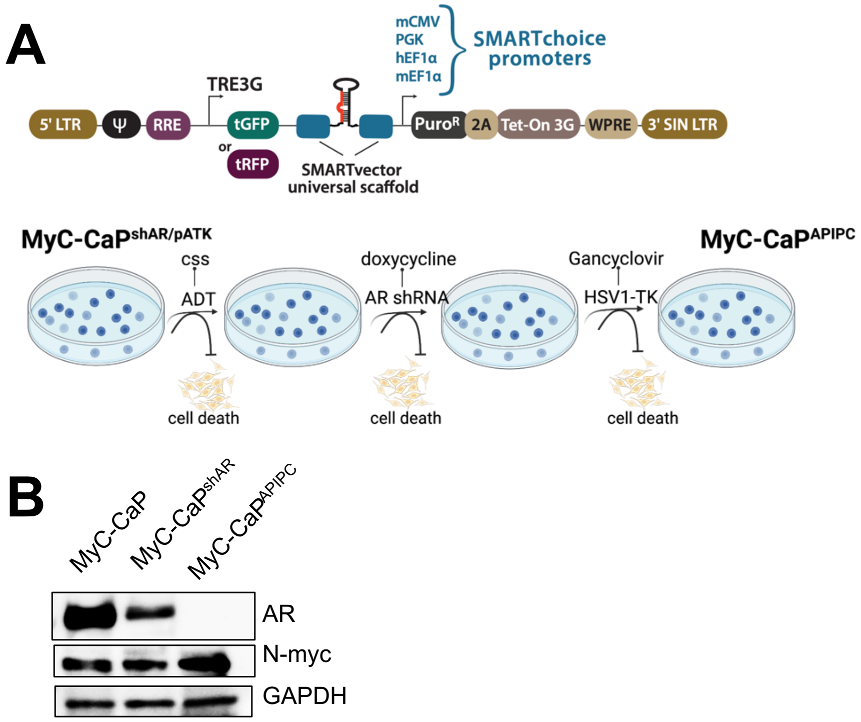

**Supp. Figure 3.** Griess test results showing nitrate concentrations in LNCaP and H660 cell lines to estimate nitrosative stress.

**Nitrate quantification (Griess test)**

- Supernatant collected when cells were confluent
- 8.5 x $10^5$ cells for lysate

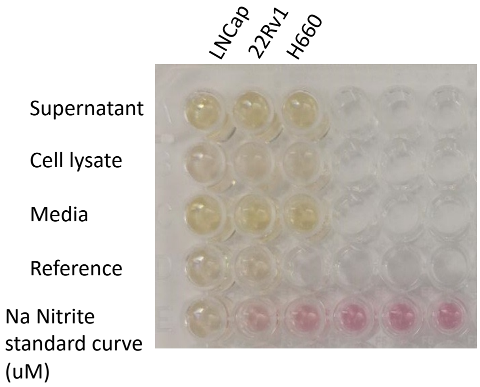


| Na Nitrite standard curve (uM) | 1 | 20 | 40 | 60 | 80 | 100 |
|---|---|---|---|---|---|---|
| Abs 548 nm | 0.097 | 0.166 | 0.231 | 0.263 | 0.282 | 0.271 |

Abs 548 nm

| | LnCap | 22Rv1 | H660 |
|---|---|---|---|
| Super | 0.107 | 0.110 | 0.099 |
| Cell lysate | 0.161 | 0.586 | 0.211 |
| Media | 0.093 | 0.097 | 0.091 |
| Reference | 0.097 | 0.097 | |

Nitrite (uM)

| | LnCap | 22Rv1 | H660 |
|---|---|---|---|
| Super | 9.7 | 6.7 | 6.3 |
| Cell lysate | 32.17 | 209.25 | 53 |

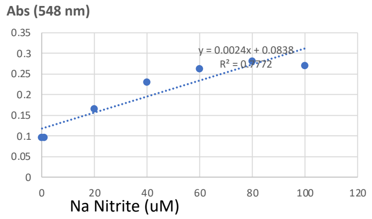

**Supp. Figure 4.** Bar graph showing overall calcium levels in 22Rv1 cells overexpressing MYCC or MYCN compared to control 22Rv1 cells. Data represent mean ± standard deviation from three independent biological replicates ($p < 0.001$, two-way ANOVA)

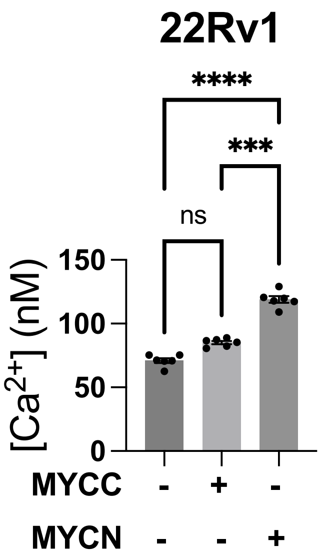

**Supp. Figure 5.** Results showing the number of mitochondria in the MyCaP, MyCaP$^{shAR}$ and MyCaP$^{APIPC}$ cells (an indicator of cellular health and bioenergetics) using quantitative fluorescence microscopy, employing MitoTracker Green FM.

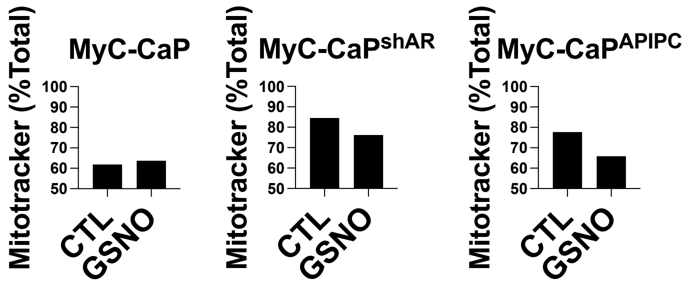

**Supp. Figure 6.** (a) showing the inhibitory effects (if any) of GSNO treatment (50uM) on the ER stress markers in DU145, PC3 and H660 cells. (b) Shows the inhibitory effects of GSNO on the colony-forming and (c) cell proliferating capabilities of MyCaP, MyCaP$^{shAR}$ and MyCaP$^{APIPC}$ cells.

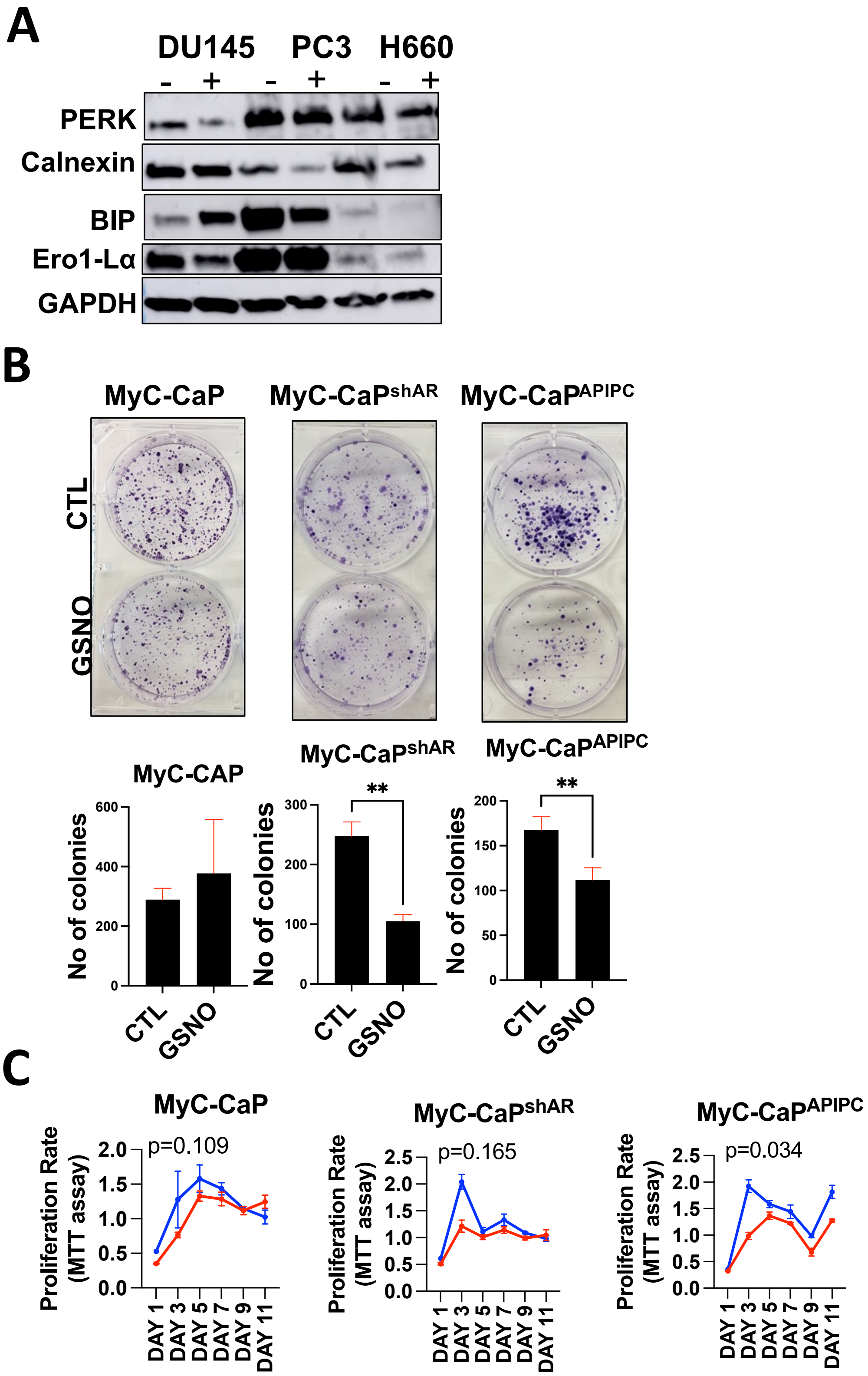

**Supp. Figure 7.** (a) shows the Gene Expression Profiling Interactive Analysis (GEPIA) of CHOP, NOS3, PDI, and MYCN, respectively, to evaluate the role of ER stress markers in distant metastasis in high-grade PCa patients.

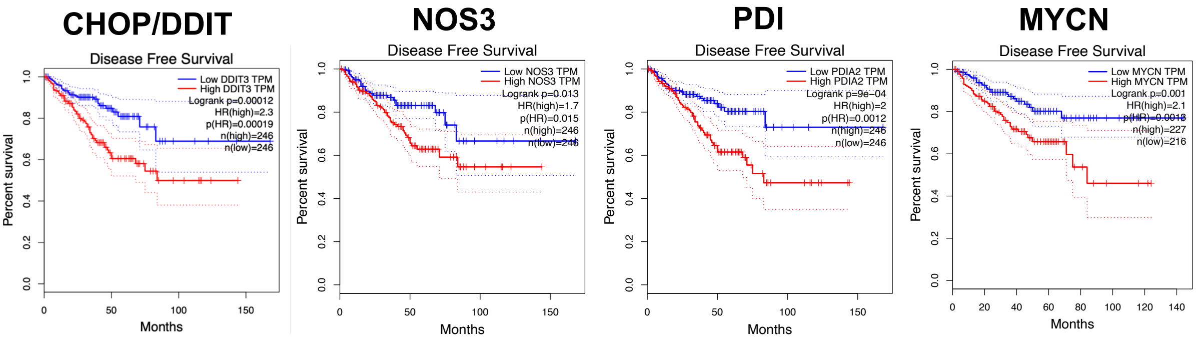

**Supp. Figure 8.** (a) Showing the outcomes of GPS-SNO 1.0 software, which identified a total of 16 cysteine residues on the MYCN protein, conforming to the acid-base nitrosylation conservative motif. Among these, Cys4, Cys186, and Cys464 showed the highest predicted thresholds for S-nitrosylation (highlighted as yellow). (b) Sanger sequencing data to confirm the site-directed mutations at Cys4, Cys186, and Cys464.

**A**

```
n-myc (https://www.uniprot.org/uniprotkb/P04198/entry)
>sp|P04198|MYCN_HUMAN N-myc proto-oncogene protein OS=Homo sapiens OX=9606
GN=MYCN PE=1 SV=2
MPSCSTSTMPGMICKNPDLEFDSLQPCFYPDEDDFYFGGPDSTPPGEDIWKKFELLPTPP
LSPSRGFAEHSSEPPSWVTEMLLENELWGSPAEEDAFGLGGLGGLTPNPVILQDCMWSGF
SAREKLERAVSEKLQHGRGPPTAGSTAQSPGAGAASPAGRGHGGAAGAGRAGAALPAELA
HPAAECVDPAVVFPFPVNKREPAPVPAAPASAPAAGPAVASGAGIAAPAGAPGVAPPRPG
GRQTSGGDHKALSTSGEDTLSDSDDEDDEEEDEEEEIDVVTVEKRRSSSNTKAVTTFTIT
VRPKNAALGPGRAQSSELILKRCLPIHQQHNYAAPSPYVESEDAPPQKKIKSEASPRPLK
SVIPPKAKSLSPRNSDSEDSERRRNHNILERQRRNDLRSSFLTLRDHVPELVKNEKAAKV
VILKKATEYVHSLQAEEHQLLLEKEKLQARQQQLLKKIEHARTC

All Sites:
4    ****MPSCSTSTMPG     4.533    0 Cluster B
14   STMPGMICKNPDLEF     1.755    0 Cluster B
27   EFDSLQPCFYPDEDD     0.372    0 Cluster A
115  NPVILQDCMWSGFSA     17.015   0 Cluster C
186  LAHPAAECVDPAVVF     3.707    0 Cluster B
323  SELILKRCLPIHQQH     0.388    0 Cluster A
464  KIEHARTC*******     3.967    0 Cluster B

Low Threshold:
4    ****MPSCSTSTMPG   4.533  2.443  Cluster B
186  LAHPAAECVDPAVVF   3.707  2.443  Cluster B
464  KIEHARTC*******   3.967  2.443  Cluster B

Medium Threshold:
4    ****MPSCSTSTMPG   4.533  2.454  Cluster B
186  LAHPAAECVDPAVVF   3.707  2.454  Cluster B
464  KIEHARTC*******   3.967  2.454  Cluster B

High Threshold:
4     ****MPSCSTSTMPG   4.533 2.443 Cluster B
186   LAHPAAECVDPAVVF   3.707 2.443 Cluster B
464   KIEHARTC*******   3.967 2.443 Cluster B
```

**B**

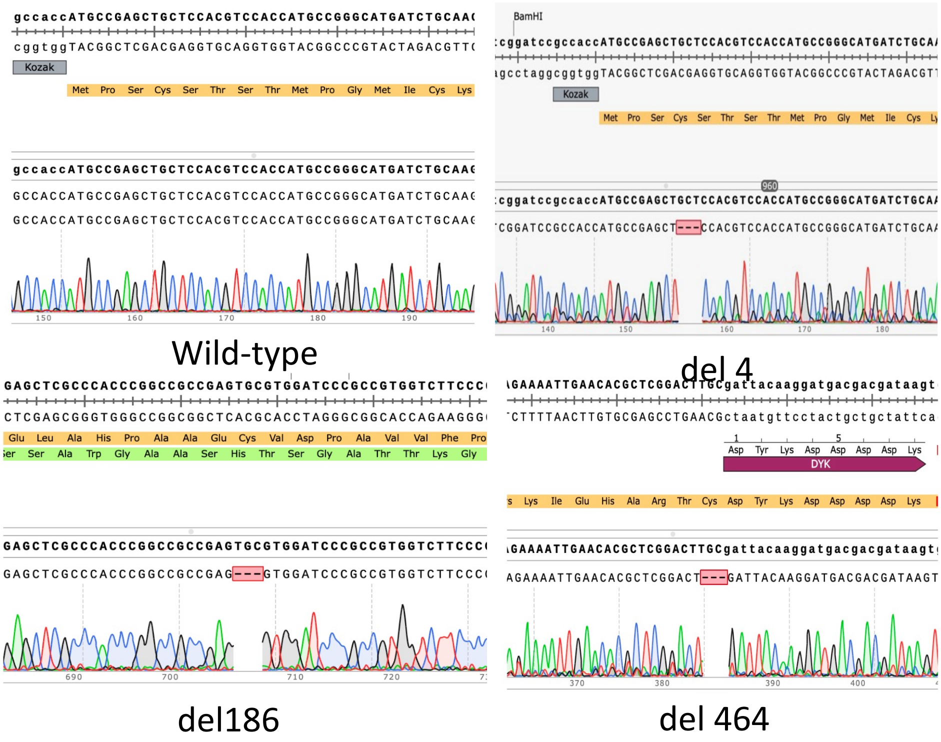

**KEY RESOURCE TABLE**

| REAGENT or RESOURCE | SOURCE | IDENTIFIER |
| --- | --- | --- |
| **Antibodies for Western Blotting** | | |
| N-Myc (D4B2Y) | Cell Signaling | 51705S |
| N-Myc | Abcam | ab24193 |
| Synaptophysin | Abcam | ab32127 |
| Chromogranin A | Abcam | ab45179 |
| BIP | Cell Signaling | 9956 |
| Calnexin | Cell Signaling | 9956 |
| Ero1-Lα | Cell Signaling | 9956 |
| IRE1α | Cell Signaling | 9956 |
| CHOP | Cell Signaling | 9956 |
| PERK | Cell Signaling | 9956 |
| PDI | Cell Signaling | 9956 |
| XBP-1S, no catalog #, was given by Himanshu's wife | | |
| P-IRE1α, no catalog #, was given by Himanshu's wife | | |
| Androgen receptor | Abcam | ab133273 |
| GAPDH | Abcam | ab181603 |
| GAPDH | Cell Signalling | sc-47724 |
| GAPDH | Cell Signalling | 2118S |
| Anti-mouse IgG | Cell Signaling | 7076S |
| Anti-rabbitt IgG | Cell Signaling | 7074S |
| **Antibodies for immunohistochemistry** | | |
| CHG | Cell signaling | 70076 |
| SYP | Abcam | ab178945 |
| N-myc | Cell signaling | 4370 |
| **Flow cytometry dye** | | |
| MitoTracker™ Dye for Flow Cytometry | Thermo Fisher Scientific | M46750 green |
| **Cell culture media** | | |

| | | |
|---|---|---|
| F-12K (PC3 cell line) | ATCC | 30-2004 |
| EMEM (DU145) | ATCC | 30-2003 |
| RPMI-1640 (H660, 22Rv1) | ATCC | 30-2001 |
| Antibiotic antimicotic solution 100X | Sigma-Aldrich | A5955-100ML |
| **Biological Samples** | | |
| Prostate Biopsies | University of Miami | |
| **Chemicals, Peptides, and Recombinant Proteins** | | |
| Tri-reagent | Sigma-Aldrich | T9424 |
| iQ™ SYBR® Green Supermix | Bio-Rad | 10296-028 |
| SYBR Universal PCR Master Mix | BioRad | 1725274 |
| RIPA Buffer | Cell Signaling | PHG0026 |
| Phosphatase Arrest™ Phosphatase Inhibitor Cocktail | G-Biosciences | 9806 |
| Protease Arrest™ Protease Inhibitor Cocktail | G-Biosciences | 786-450 |
| Pierce™ ECL Western Blotting Super Signal Pico Plus | Thermo Scientific | 34580 |
| ABC detection kit | Abcam | ab64264 |
| S-Nitrosylated Protein Detection Kit | Cayman Chemical | 10006518 |
| MTT reagent | Sigma Aldrich | CT01 |
| Signal-SeekerTM Ubiquitination Detection Kit | Cytoskeleton | BK161 |
| Griess Reagent Kit for Nitrite Determination | Molecular probes | G-7921 |
| **Critical Commercial Assays** | | |
| iScript cDNA Synthesis Kit | Bio-Rad | |

| **Software and Algorithms** | | |
|---|---|---|
| Adobe Illustrator | Adobe Systems, San Jose, CA | https://www.adobe.com/ca/products/illustrator.html |

| | | |
|---|---|---|
| FlowJo software V10 | FlowJo, LLC | https://www.flowjo.com/ |

### MATERIAL & METHODS:

#### Cell Culture

PC3 (CRL-2505), DU145 (CRL-1740) and H660 (CRL-5813) were purchased from ATCC. PC3 cells were maintained in F12K medium. DU145 cells were grown in DMEM medium. 22Rv1 cell line was cultured in RPMI-1640. NCI-H660 (CRL-5813) were cultured in HITES medium. All cells but H660 (5%) were cultured in presence of 10% fetal bovine serum (Gibco), 100 U/ml penicillin, and 100 µg/ml streptomycin.

MyC-CaP cells were stably transfected with SMART shRNA doxycycline-inducible Lentiviral vector (Horizon Discovery Ltd. ) to generate MyC-CaP$^{shAR}$ cell lines. These cells were further modified via Lipofectamine 2000 (Invitrogen) transfection of a plasmid encoding a triple-probasin-driven herpes thymidine kinase (HSV-TK) and a puromycin resistance cassette, which was a gift from Peter Nelson's lab. A clonal population of this cell line derived from puromycin (Invitrogen) selection and serial dilution in a 96-well plate, which we refer to as MyC-CaP $^{shAR/pATK}$, was subjected to total AR pathways suppression (TAPS): for two weeks the cells were grown in RPMI1640+10%CSS; at week 3, media was supplemented with 1 mg/mL doxycycline. Media was changed every 3–4 days and MyC-CaP$^{shAR/pATK}$ was maintained under TAPS for five months. A surviving colony of proliferating cells emerged. Following a 3-month expansion, this population of cells was treated with 50 µM ganciclovir (GCV; InvivoGen) for two weeks to eliminate any cells still robustly expressing an AR transcriptional program. We referred to the surviving population as MyC-CaP $^{APIPC}$.

#### MTT assay

PC3, DU145 and H660 cells were seeded in 96-wells plate in quadruplets and were treated with varying doses of GSNO. MTT (3-(4,5-dimethylthiazole)-2,5-diphenyltetrazolium bromide, Sigma Aldrich CT01) assay reagents were added, and the absorbance was measured at 562 nm after 0, 3, 5, 7, and 9 days.

For RNA/protein collection, when cells were around 80% confluent, growth media was changed by those using charcoal stripped FBS. Then, treated with GSNO 50uM for different time points. After treatment, a colony formation assay using 2000 cells/well was done and stained with Cristal Violet, and cells were collected for RNA/protein isolation.

**Preparation of RNA and quantitative real-time PCR**

Total RNA was extracted from cells using the Trireagent (Sigma-Aldrich T9424) method and then reverse transcribed to complementary DNA using High-Capacity cDNA Reverse Transcription Kits (Applied Biosystems, USA) according to the manufacturer's protocol. The quantitative RT-PCR for indicated genes (MYCN, CHGA, SYP, ENO2, NCAM1, Ki67) was performed in SYBR Universal PCR Master Mix (BioRad 1725274). Quantitation of mRNAs was performed using BIORAD™ Gene Expression Assays according to the manufacturer's protocol. Samples were analyzed using the BIORAD sequence detection system. All PCRs were performed in triplicate, and the specificity of the reaction was determined by melting curve analysis at the dissociation stage. The relative quantitative method was used for the quantitative analysis. The calibrator was the average ΔCt from the untreated cells. The endogenous control was glyceraldehyde 3-phosphate dehydrogenase (GAPDH).

**Western blotting**

Cells were harvested and lysed in RIPA lysis buffer containing protease and phosphatase Inhibitor Cocktail (Sigma, St. Louis, MO, USA). Protein expression was studied by exposing the membranes to antibodies against NEPC markers CHG (Abcam, ab45179), SYP (ab32127), DOT1L (), EZH2 (Cell Signalling), MYCN (Cell Signalling, 51705 or Abcam ab24193), ER stress markers kit (BIP, Calnexin, Ero1-Lα, IRE1α, CHOP, PERK, PDI. Cell Signaling 9956), XBP-1S, P-IRE1α, androgen receptor (ab133273) and GAPDH (Santa Cruz Biotechnology, sc-47724, Cell Signalling 2118S). Immunoreactive bands were visualized using the Chemiluminescent Super Signal Pico Plus Kit (Thermo Scientific 34580). Protein bands were quantified using ImageJ software and normalized against GAPDH.

**Immunohistochemistry**

For immunohistochemistry, tissue sections were stained with hematoxylin and eosin and analyzed by a genitourinary pathologist. For immune-histochemical staining, tumor xenograft tissues were fixed in 10% buffered formalin and embedded in paraffin. 5 µm thick sections were deparaffinized and rehydrated in sequential xylene and graded ethanol. Antigen retrieval was performed in 10 mM citrate buffer (pH 6.0) in a microwave oven. Peroxidase and non-specific protein blocking were done as per the instructions using the Abcam ABC detection kit (ab64264) and incubated with the following primary antibody dilutions: CHG (cell signaling, 70076), SYP (Abcam, ab178945) and MYCN (Cell signaling, 4370) with 1:150 dilution. They were subsequently incubated with biotinylated goat anti-polyvalent secondary antibody, followed by development using DAB substrate as per the instructions on the kit. All sections were lightly counterstained with hematoxylin and mounted with Cytoseal XYL. Images were taken using a brightfield microscope (Nikon E200) at 10X and 40X magnification, and quantification was done using ImageJ 1 Front Endocrinol (Lausanne).

#### Agonistic and antagonistic assays

22Rv1 cells were stably transduced with MYC viruses (PLV-10005, Cellomics) at a concentration of $1x10^8$ TU/ml. After addition of viruses, we waited for 48 hours for cells to be infected. Puromycin selection antibiotic was used at a concentration of 5ug/ml to select effectively transduced cells. Post selection, the remaining cells were allowed to grow further to at least 3 passages to use it for any further experiments.

#### S-Nitrosylation assay

We evaluated if the post-translational modification S-nitrosylation, induced by GSNO, is acting as a degradation signal for MYCN. To do that we performed the biotin-switch assay following the manufacturer’s guidelines (S-Nitrosylated Protein Detection Kit (Biotin Switch), Item No. 10006518, Cayman Chemical, Ann Arbor, MI, USA). A small piece of tumor tissue or cell pellet was taken and washed twice with Wash Buffer. The pellets were resuspended in “Buffer A containing Blocking Reagent” and incubated for 30 min at 4 °C with shaking. The incubated samples were centrifuged, and the supernatant was transferred to 15 ml centrifuge tubes. Two milliliters of ice-cold acetone were added to each sample, and the mixture was incubated at −20 °C for at least 1 h. The protein from each sample was pelleted by centrifugation. “Buffer B containing Reducing and Labeling Reagents” was added to resuspend the proteins, with incubation for 1 h at room temperature. The biotinylated protein was precipitated by acetone and rehydrated with the appropriate amount of Wash Buffer. A total of 50 µg of protein was used for labeling and running the standard western blot for detecting the nitrosylated protein.

Besides, we deleted the S-nitrosylated sites in this protein by site directed mutagenesis. MYCN ORF clone was procured from GenScript Biotech (NJ, USA). Site-directed mutagenic changes were performed to incorporate 3 cysteine deletions at C4, C186, and C464 (as determined by GPS-SNO 1.0 software with a high threshold) using Quick Change Lightning Multi Site-Directed Mutagenesis Kit (Agilent Technologies, USA), as previously described. Briefly, 40 ng plasmid was subjected to PCR amplification as per standard kit guidelines using mutagenic primers designed specifically for cysteine deletions at these specific sites. Following PCR, 10 µl of the product was subject to DpnI digestion for 5 min at 37 °C and transformed into chemically competent DH5α cells (NEB, USA) by heat shock at 42 °C for 30 sec. The resulting transformants were grown in SOC media for 1 hr at 37 °C and selected overnight on LB agar plates containing 100 µg/ml of Ampicillin. The following day, single colonies were selected and further grown. Plasmid isolation and purification were done using a plasmid miniprep kit (Qiagen, Germany) as per standard instructions. Sanger sequencing for confirmation of deletion was done by Genewiz, USA. For verification of these deletions, the confirmed wild-type, as well as mutant clones, were transfected in NCI-H660 cells using Lipofectamine 3000 reagent. After transfection, cells were treated with/without 50 µM GSNO, and cells were collected after 48 h to do western blots using specific antibodies for ER stress.

**Animals**

The animal protocol was approved by the Institutional Animal Care and Use Committee of the University of Miami Miller School of Medicine, Miami, FL. SCID (3 weeks old) mice were purchased from Jackson Laboratories. Castration experiments were performed in all the mice. For castration, mice were anesthetized using Isoflurane (Abbott Laboratories). The perineal region was cleaned with ethanol and a betadine scrub (VWR, AJ159778), and sterile dissecting shears were used to make a 4–5 mm incision. Using two sterile forceps, the testes were located, and a ligature was made around the testicular vessels and the tunica albuginea that encases the testes. The testes were amputated with dissecting shears, and the scrotum was sutured closed with 6-0 Ethicon black monofilament nylon (Ethicon Inc., 1665). A local triple antibiotic was applied over the region of the wound to facilitate healing. SCID mice were grouped into control and experimental groups. All the mice were grafted with 1 million NCI-H660 cells orthotopically. The mice were distributed randomly into the control and experimental group. The experimental group received 10 mg/kg/day of GSNO treatment intraperitoneally (IP) 3 times/week for 2 weeks, while the control group received PBS IP. Each group had 10 mice each. After treatment, animals survived for an additional two weeks before humanely sacrificing them. At the end time point, blood was collected via cardiac puncture, and tumor grafts, lungs, and spleen were harvested for further analysis. Tumor volume (V) was measured regularly (blinded) until the mice were sacrificed by measuring the length (L) and width (W) of the tumor with calipers by using the formula: $V = 1/2(\text{length} \times \text{width}^2)$. Tumor growth was also monitored by IVIS.

**IP-Mass Spectrometry Analysis**

Tissue lysates were co-immunoprecipitated (co-IP) using MYCN antibody using Thermo Scientific™ Pierce™ Classic Magnetic IP/Co-IP Kit as per the standard guidelines mentioned in the kit. The MYCN ab was first added to the sample to form an immune complex that is then bound to the magnetic beads. The complex was washed to remove non-bound material and a low-pH elution buffer dissociated the bound immune complex from the Protein A/G. The beads were removed from the solution manually using a magnetic stand.

The eluted peptide mixtures were analyzed using a nanoflow liquid chromatograph (U3000, Dionex, Sunnyvale, CA) coupled to an electrospray ion trap mass spectrometer (LTQ-Orbitrap, Thermo, San Jose, CA) in a data-dependent manner for tandem mass spectrometry peptide sequencing experiments. Both MASCOT and SEQUEST search results were summarized in Scaffold 2.0. The integrated peak areas for phosphotyrosine peptide quantification were calculated from extracted ion chromatograms (EIC) using QuanBrowser from Xcalibur 2.0.

**GSNO reductase (GSNOR) activity assay**

Cell lines were washed, trypsinized and homogenized by sonication (30’ in ice) in a solution containing 20mM Tris-HCl (pH 8.0), 0.5 mM EDTA, 0.1% NP-40 and 1mM phenylmethylsulphonyl fluoride (PMSF). To detect GSNO reductase enzymatic activity, 80ug total protein was incubated with reaction buffer (20mM Tris-HCl, pH 8.0, 0.5mM EDTA) with 0 (negative control) or 200 uM NADH in the presence of GSNO 50 uM. Then, GSNO reductase activity was measured by reading absorbance at 340 nm every 5’ during 30’ (Nature, 410, 490–494, Liu 2001).

**Calcium Release Estimation**

To assess the impact of nitric oxide (NO) supplementation on ER stress-induced calcium release in prostate cancer cell lines, we used a fluorescence-based approach. LNCaP (primary prostate adenocarcinoma), 22Rv1 (castration-resistant prostate cancer), and H660 (neuroendocrine prostate cancer) cells were cultured in RPMI 1640 media supplemented with 10% FBS and 1%

penicillin-streptomycin at 37°C with 5% CO2. Cells grown on a monolayer and treated with either 50 μM S-nitroso glutathione (GSNO) or vehicle, were loaded with 2.5 μM Fura-2 AM (Molecular Probes, Eugene, OR), a dual excitation calcium-sensitive dye, for 20 minutes at room temperature, followed by a 30-minute de-esterification period in the dark. Then, fluorescence (Ex: 340/380 nm, Em: 515 nm) was recorded in an IonOptix spectrofluorometer (IonOptix LLC, Westwood, MA). The calibration was performed on the cells acquiring fluorescence data in a $Ca^{2+}$-free and then a $Ca^{2+}$-saturating (5 mmol/L) solutions, both containing 10 μmol/L ionomycin (Sigma, St. Louis, MO) until reaching a minimal ($R_{min}$) or a maximal ($R_{max}$) ratio value, respectively. Intracellular $Ca^{2+}$ ($[Ca^{2+}]i$) was calculated using the following equation:

$$[\boldsymbol{Ca^{2+}}]_{\boldsymbol{i}} = K_d \times \frac{S_{f2}}{S_{b2}} \times \frac{(R - R_{\min})}{(R_{max} - R)}$$

$K_d$ (dissociation constant) in adult myocytes was taken as 224 nmol/L. The scaling factors $S_{f2}$ and $S_{b2}$ were extracted from calibration as described by Dulce et al [65].

**Measurement of Oxygen consumption rate and Extracellular acidification rate**

Oxygen consumption rate (OCR) and Extracellular acidification rate (ECAR) were measured using the Seahorse XF Pro Analyzer (Agilent Technologies). Mitochondrial stress test was performed to measure OCR per manufacturer's instructions. Briefly, $4x10^4$ H660 or $3x10^4$ LNCaP cells were plated in complete growth media into each well of a 96-well Seahorse microplate and incubated overnight in 5% $CO_2$ at 37ºC. Cells were then treated with GSNO for 48 hours. Following GSNO treatment, cells were washed twice, incubated (in non-$CO_2$ incubator at 37ºC for 1 hour), and analyzed in XF assay media (non-buffered RPMI containing 10 mM glucose, 2 mM L-glutamine, and 1 mM sodium pyruvate, pH 7.4) at 37ºC, under basal conditions and in response to 2 μM oligomycin (Sigma), 2 μM fluoro-carbonyl cyanide phenylhydrazone (FCCP) (Sigma) and 0.5 μM rotenone (Sigma)/0.5 μM antimycin A (Sigma). Data were analyzed by the Seahorse XF Cell Mito Stress Test Report Generator. OCR (pmol $O_2$/min) values were normalized to the protein content.

ECAR values were measured by performing glycolysis stress test according to manufacturer's instructions. Briefly, $4x10^4$ H660 or $3x10^4$ LNCaP cells were plated in complete growth media into each well of a 96-well Seahorse microplate and incubated overnight in 5% $CO_2$ at 37ºC. Cells were then treated with GSNO for 48 hours. Following GSNO treatment, cells were washed twice, incubated (in non-$CO_2$ incubator at 37ºC for 1 hour), and analyzed in XF assay media (non-buffered RPMI containing 2 mM L-glutamine, pH 7.4) at 37ºC, under basal conditions and in response to 10 mM glucose (Sigma), 2 μM oligomycin (Sigma), and 50 mM 2-deoxy-D-glucose (Sigma). Data were analyzed by the Seahorse XF Cell Glycolysis Stress Test Report Generator. ECAR (mpH/min) values were normalized to the protein content.

**Nitrite Determination (Griess Kit, Molecular Probes G7921)**

Nitrite levels were quantified using the Griess Reagent Kit (Molecular Probes, G7921) according to the manufacturer's instructions. Briefly, supernatant was collected from T75 flasks containing cells grown to confluency. The cells were cultured in the appropriate media, and once confluency was reached, the supernatant was harvested for analysis.

To measure nitrite, 150 μL of the collected supernatant was mixed with 150 μL of the Griess reagent in a 96-well plate, in triplicate, for each sample. A standard curve was prepared using known concentrations of sodium nitrite ranging from 0 to 100 μM in the same 96-well format. The plate was incubated at room temperature for 30 minutes, protected from light, allowing the color to develop.

Absorbance was measured at 548 nm using a microplate reader (insert model here), and the nitrite concentration in the samples was calculated by comparing the absorbance values to the standard curve. All measurements were performed in triplicate, and results were expressed as mean ± standard deviation.

**Transcriptomic Data Analysis and Survival Correlation**

RNA-seq data from 500 patients in the TCGA PRAD cohort and 4,983 patients from the Decipher GRID database were analyzed to investigate gene expression changes associated with PCa progression. Patients were stratified based on clinical parameters including Gleason scores (GS), age, PSA levels, and therapy history. The RNA-seq data were obtained in raw count format. Initial preprocessing involved normalization using the transcripts per million (TPM) method to account for differences in sequencing depth across samples. Variance-stabilizing transformation (VST) was applied to the normalized data to reduce heteroscedasticity and improve interpretability for downstream statistical analyses. Differential expression analysis was conducted using the DESeq2 package (version 1.26.0) in R. Raw read counts were modeled with a negative binomial distribution to estimate fold changes in gene expression between patient groups. Comparisons were made between patient subgroups based on Gleason scores and other clinical characteristics. The results of the differential expression analysis were filtered using a false discovery rate (FDR) of less than 0.05 to control for multiple testing, and only genes with a log2 fold change of ±1 or higher were considered for further analysis. Correlation analysis was performed to assess the relationships between gene expression profiles. Pearson correlation coefficients were calculated for selected genes across the Decipher GRID dataset. Prior to correlation analysis, the data were mean-centered and scaled. A two-tailed t-test was used to evaluate the statistical significance of the correlation coefficients, with p-values less than 0.05 considered significant. Correlation matrices were visualized using heatmaps generated with the ggplot2 package in R. Survival analysis was carried out to assess the clinical relevance of gene expression levels using the cBioPortal platform. Kaplan-Meier survival curves were generated to compare overall survival (OS) between patient groups stratified by gene expression levels (high versus low), which were determined based on the median expression of each gene. Hazard ratios (HR) with 95% confidence intervals were calculated using the log-rank test to determine the strength of the association between gene expression and survival outcomes.

**Statistical Analysis**

All statistical analyses were conducted using R (version 4.0.3) and associated packages, unless otherwise specified. Data are presented as mean ± standard deviation (SD) or mean ± standard error of the mean (SEM), depending on the experiment. For comparisons between two groups, statistical significance was assessed using unpaired two-tailed Student's t-tests. For comparisons involving more than two groups, one-way or two-way analysis of variance (ANOVA) was employed, followed by post hoc Tukey's or Bonferroni correction for multiple comparisons where applicable. P-values less than 0.05 were considered statistically significant. All experiments were performed with at least three independent biological replicates unless otherwise indicated. Statistical analyses were performed using R or GraphPad Prism (version 9.0), and figures were generated using appropriate visualization tools such as ggplot2 and GraphPad Prism. A p-value of $<0.05$ was considered statistically significant across all analyses.
